# Coenzyme-Protein Interactions since Early Life

**DOI:** 10.1101/2023.10.28.563965

**Authors:** Alma Carolina Sanchez-Rocha, Mikhail Makarov, Lukáš Pravda, Marian Novotný, Klára Hlouchová

## Abstract

Recent findings in protein evolution and peptide prebiotic plausibility have been setting the stage for reconsidering the role of peptides in the early stages of life’s origin. Ancient protein families have been found to share common themes and proteins reduced in composition to prebiotically plausible amino acids have been reported capable of structure formation and key functions, such as binding to RNA. While this may suggest peptide relevance in early life, their functional repertoire when composed of a limited number of early residues (missing some of the most sophisticated functional groups of today’s alphabet) has been debated.

Cofactors enrich the functional scope of about half of extant enzymes but whether they could also bind to peptides lacking the evolutionary late amino acids remains speculative. The aim of this study was to resolve the early peptide propensity to bind organic cofactors by analysis of protein-coenzyme interactions across the Protein Data Bank (PDB). We find that the prebiotically plausible amino acids are more abundant in the binding sites of the most ancient coenzymes and that such interactions rely more frequently on the involvement of the protein backbone atoms and metal ion cofactors. Moreover, we have identified a few select examples in today’s enzymes where coenzyme binding is supported solely by prebiotically available amino acids. These results imply the plausibility of a coenzyme-peptide functional collaboration preceding the establishment of the Central Dogma and full protein alphabet evolution.

## Introduction

Organic and inorganic cofactors occupy about half of all known protein structures, expanding across all the enzyme E.C. classes (Putignano, 2018; Mukhopadhyay et al., 2019). While their role in current life is indisputable, some of the cofactors were present and apparently crucial also during life’s early evolution (Chu and Zhang 2020; Goldman and Kacar 2021; Kirschning et al., 2021; Fried et al., 2022; Kirschning, 2022). The significance of metal ions has been broadly discussed, regardless of the different origins-of-life scenarios and has somewhat overshadowed that of organic cofactors (e.g. Wächtershäuser, 1992; Russell and Hall, 1997; Lane and Martin, 2012 ; Chu and Zhang, 2020; Fried et al., 2022).

Diverse lines of evidence have however indicated that many of the extant organic cofactors (coenzymes) date back to the earliest life while their core chemistries have been detected in abiotic material such as recently reported by the Hayabusa2 mission (Holliday et al., 2007; Fried et al., 2022; Naraoka et al., 2023). At the same time, these ancient coenzymes – often of nucleotide origin– have been traced to the most ancient protein folds (such as P-loop NTPases, TIM beta/alpha-barrels, OB and Rossmann folds) that date before the Last Universal Common Ancestor (LUCA) (Goldman and Kacar, 2021; Caetano-Anollés et al., 2007; Goldman et al., 2013; Longo et al., 2020 (a); Kessel and Ben-Tal, 2022). Within the most ancient folds, tens of peptide fragments/themes have been identified throughout seemingly unrelated structural domains, and frequently found to mediate ligand binding (Söding and Lupas, 2003; Alva et al., 2015; Narunsky et al., 2020; Kolodny et al., 2021). Such themes may well represent the remnants of protoenzymes in a peptide-nucleotide world (Fried et al., 2022). Chu and Zhang recently proposed that cofactors could initially “select” the earliest primitive proteins from the vast sequence space by the ability to bind them (Chu and Zhang, 2020). More generally, binding of cofactors to peptides could thus determine the evolution of both protoenzyme function and folding preferences (Tokuriki and Tawfik, 2009).

Prior to the fixation of Central Dogma and ribosomal synthesis, peptides would condense from amino acids (or their alternatives) prebiotically abundant in the environment (Frenkel-Pinter et al. 2019; Frenkel-Pinter et al. 2020; Fried et al., 2022). Independent meta-analyses of the amino acid alphabet evolution based on different possible sources of organic material and different disciplines point towards an “early alphabet” of ∼10 residues (Ala, Asp, Glu, Gly, Ile, Leu, Pro, Ser, Thr and Val) (Higgs and Pudritz, 2009; Trifonov, 2000; Cleaves, 2010). These could be supplemented by other prebiotically plausible non-canonical amino acids while the other half of the canonical alphabet is assumed to be the product of later biosynthesis (Wong and Bronskill, 1979; Weber and Miller, 1981; Burton et al., 2012; Zaia et al., 2008). Typically, the early amino acids (canonical as well as non-canonical) are smaller and less complex, missing e.g. sulfur groups and aromatics. Additionally, the canonical early alphabet lacks positively charged residues. An emerging question therefore is whether coenzymes could bind to small proteins of prebiotic relevance and whether they could be bound by the prebiotically available residues. In such a scenario, cofactors would provide a palette of functional groups to the early peptide world which would nominate them relatively sophisticated structural and catalytic hubs (Milner-White and Russell, 2011). Alternatively, if coenzymes could not be bound by these simple amino acids, this would suggest that their pairing with peptide molecules would become relevant only after the evolution of the full amino acid alphabet.

Work from our group and others has recently demonstrated that in select cases, protein sequences re-engineered from the early amino acids can still bind to nucleic acid and nucleotide-based cofactors (Longo et al., 2020 (b); Makarov et al., 2021; Giacobelli et al., 2022). Whether this phenomenon is still seen in today’s biology, its abundance and laws represent open questions. Here, we present a systematic survey of coenzyme binding throughout the PDB database. The outcomes of our study support that the coenzyme binding characteristics by amino acids differ by their evolutionary age. Early amino acids are enriched in binding pockets of the most ancient coenzymes and the interaction relies predominantly on the protein backbone groups. Selected examples show that unlike evolutionary younger cofactors, the ancient cofactors can still be bound in proteins only by early amino acids. Our analysis therefore points to an early peptide-coenzyme significance, preceding evolution of proteosynthesis and fixation of the Central Dogma.

## Results

### Identification of coenzymes in PDB

We identified all the available structures from the PDB that interact with the 27 coenzyme classes as defined in Fischer et al., 2010. In addition, ATP (that was not included in that study) was included here. Using these parameters, we found 25,822 protein structures and 81 nucleic acid macromolecules (Supplementary Table 1, Supplementary File 1). The protein structures were assigned to 8194 UniProt (The UniProt Consortium, 2023) codes. Those UniProt sequences were clustered by 90% identity and resulted in 7399 unique UniProt entries, corresponding to 21,317 protein structures (Fig. 1). In parallel, the clustering was also performed for 30% sequence identity, resulting in 3544 UniProt codes and 9645 PDB structures.

**Fig 1:**
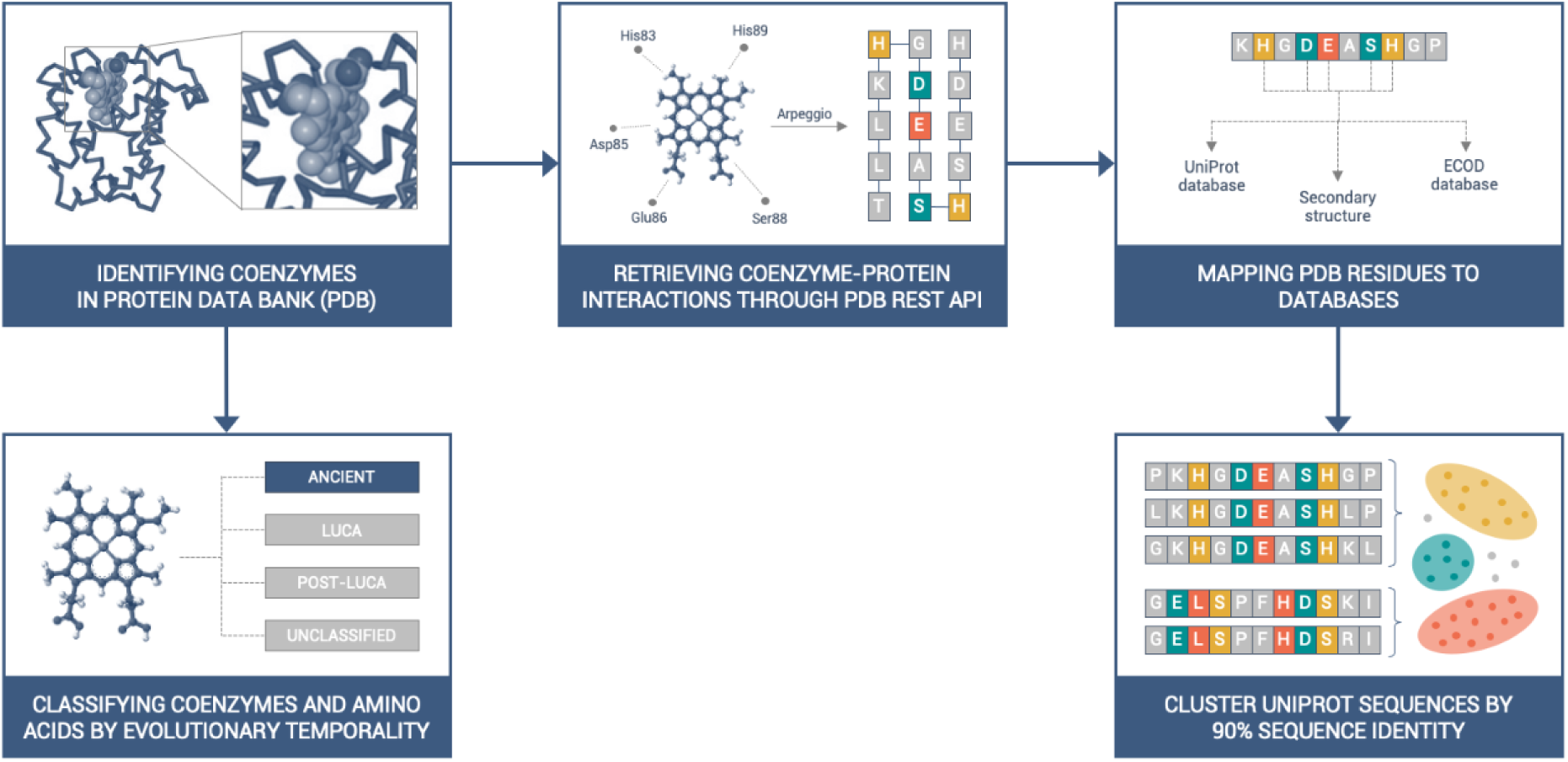
Workflow of this study. All available coenzymes in the PDB were identified according to the CoFactor database (Fischer et al., 2010). The PDB entries of structures bound to coenzymes were downloaded programmatically through the PDBe REST API (pdbe.org/api), including the interatomic cofactor-protein interactions, calculated by Arpeggio (Jubb et al, 2017). The coenzyme binding amino acids were mapped to Uniprot databases via SIFTS (Velankar et al., 2013; Dana et al., 2019). PDB entries were grouped by UniProt code; redundancy was removed by clustering the UniProt sequences by 90% (and in parallel also 30%) sequence identity.

The interaction ratio method was adopted to identify the most relevant residues in coenzyme binding sites. For each protein (unique UniProt ID) we defined the cofactor binding site as a subset of amino acids that appeared to interact with the cofactor in at least 50% of the structures of that particular protein within our dataset to pinpoint the amino acids that are important for the interaction. This methodology does not consider any qualitative criteria (e.g. resolution, R-factor, Clashscore).

Our database is composed of protein structures from members of the three cellular domains – Bacteria (54.3%), Archaea (6.2%), Eukaryota (37.8%) – as well as Viruses (1.5%), metagenomes and unassigned entries (0.3%) (Supplementary Table 1).

### Evolutionary classification of coenzymes

To differentiate the evolutionary age of the analyzed coenzymes, we further adapted the classification system from Fried et al., 2022. This system encompasses four primary categories (or temporalities) and one additional subcategory: i) "Ancient" coenzymes, including the subcategory "Nucleotide derived"; ii) "LUCA" coenzymes; iii) "Post-LUCA" coenzymes; and iv) "Unclassified" coenzymes (Fig. 2).

**Fig 2:**
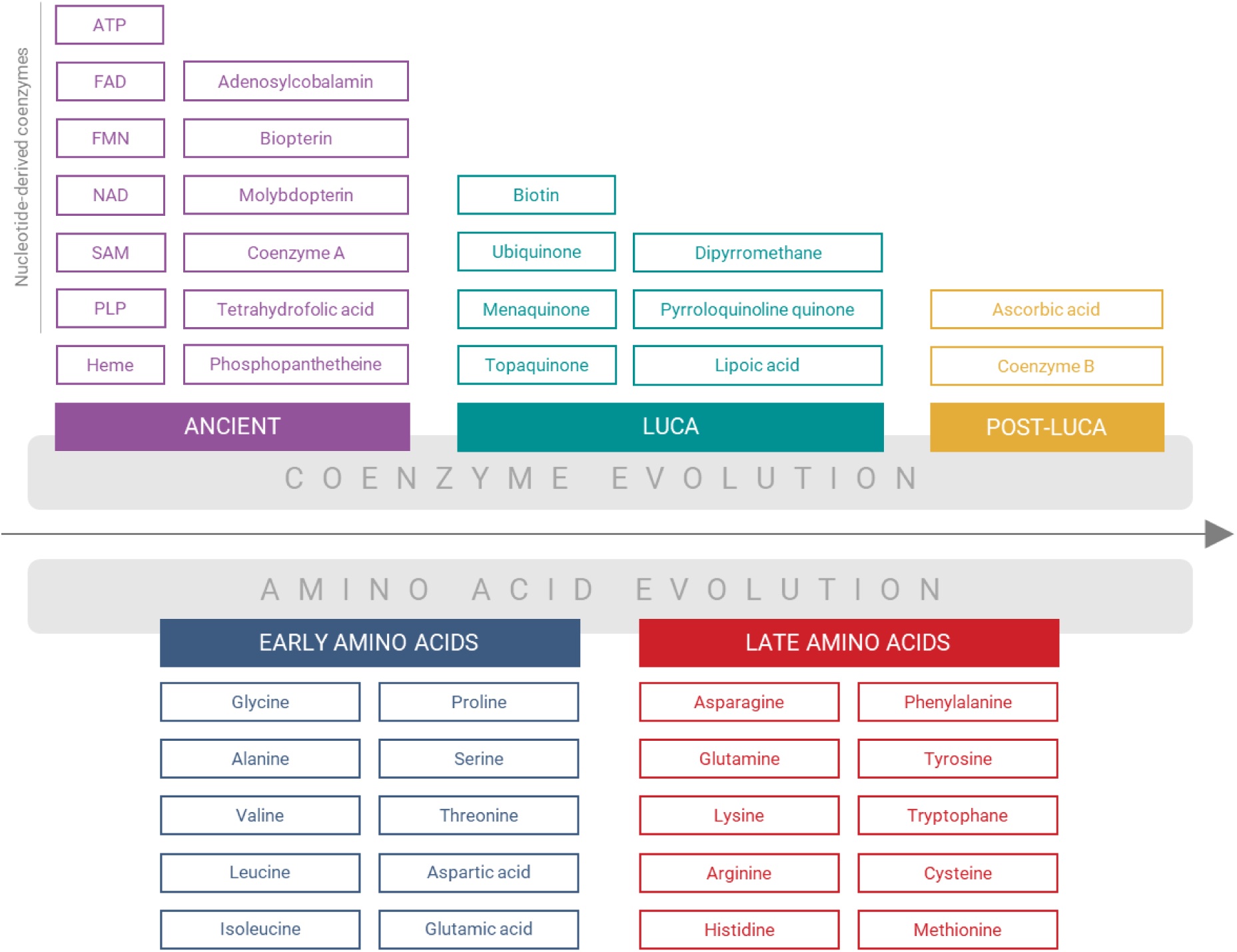
Classification of coenzymes and amino acids by their assumed evolutionary temporality. The “Unclassified” coenzymes Thiamine diphosphate, Coenzyme M, Factor F430 and Glutathione are not shown in the scheme.

“Ancient” coenzymes comprise those that could be prebiotically synthesized, according to available studies (Miller and Schlesinger, 1993; Keefe et al., 1995; Holliday et al., 2007; Kirschning, 2021; Menor-Salván et al., 2022; Pinna et al., 2022); while the subcategory "Nucleotide derived" includes cofactors chemically derived from nucleotides (White, 1975; Monteverde et al., 2017). “LUCA” coenzymes were presumably present in the last universal common ancestor (LUCA) and exhibit a universal distribution among Bacteria, Archaea, and Eukarya, although their prebiotically feasible synthesis was not established. “Post-LUCA” coenzymes likely originated only after the divergence of the three cellular domains, mirrored in their non-universal distribution. “Unclassified” coenzymes do not conform to the classification scheme. As a typical representative of the latest category, Coenzyme M has been synthesized under prebiotic conditions (Miller and Schlesinger, 1993; Kirschning, 2021), nonetheless, its biosynthetic pathways in Archaea and Bacteria have been shown to arise through convergent evolution and it is mainly prevalent in methanogens (Wu et al., 2022). Factor F430 is a coenzyme only distributed in methanogens (Thauer and Bonacker, 1994), although its precursors have been synthesized prebiotically (Seitz et al., 2021). Glutathione is another example of a coenzyme with restricted biological distribution, being mainly in eukaryotes, Gram-negative bacteria, and one archaea phylum (Copley and Dhillon, 2002) and the feasibility of its prebiotic synthesis remains unclear (Bonfio et al., 2017). Thiamine diphosphate was also designated as Unclassified. Although the definitive prebiotic synthesis of Thiamine diphosphate remains unclear, preliminary investigations conducted by Aylward (2006) and Aylward & Bofinger (2006) suggest its presence in the prebiotic world. Its nucleotide nature (White, 1975) and the existence of its universal riboswitch (Barrick and Breaker, 2007) provide compelling evidence of its potential status as an Ancient coenzyme.

The Ancient coenzymes represent the most abundant temporality of our PDB dataset, dominated by ATP, NAD, Heme, FAD, SAM, and CoA structures and amounting to 94% of all analyzed structures in our database grouped by UniProt codes. Within the enzyme E.C. classification, oxidoreductases and transferases represent the classes with the most abundant coenzyme content. While the LUCA, Post-LUCA and Unclassified coenzymes are typically found in specific enzyme classes, the Ancient coenzymes are distributed across all the E.C. classes (Supplementary Fig. 1).

### Distribution of amino acids in the coenzyme binding sites

We hypothesized that the evolutionary significance of individual coenzyme classes would be reflected in distinct amino acid binding propensities as a smaller “early” protein alphabet apparently preceded its canonical version. The abundance of residues that compose each coenzyme binding site was analyzed and examined with respect to the order by which individual amino acids have been reported to enter the protein canonical alphabet (Higgs and Pudritz 2009) (Fig. 3; Supplementary Table 2). The binding site composition for both 90% and 30% identity datasets revealed that the occupancy of early amino acids is higher in Ancient coenzyme binding sites and tends to decrease in LUCA and Post-LUCA cofactor sites. Overall, for the 90% dataset the average occupancy of early vs. late amino acids in the Ancient sites is 61% vs 39% while this ratio decreases to 53% vs 47% for the LUCA and 47% vs 53% in Post-LUCA sites. These numbers follow the same trend for the 30% identity dataset and throughout the rest of the analysis, the 90% identity dataset – which includes a higher number of proteins – was evaluated for more robust statistical analysis.

**Fig 3:**
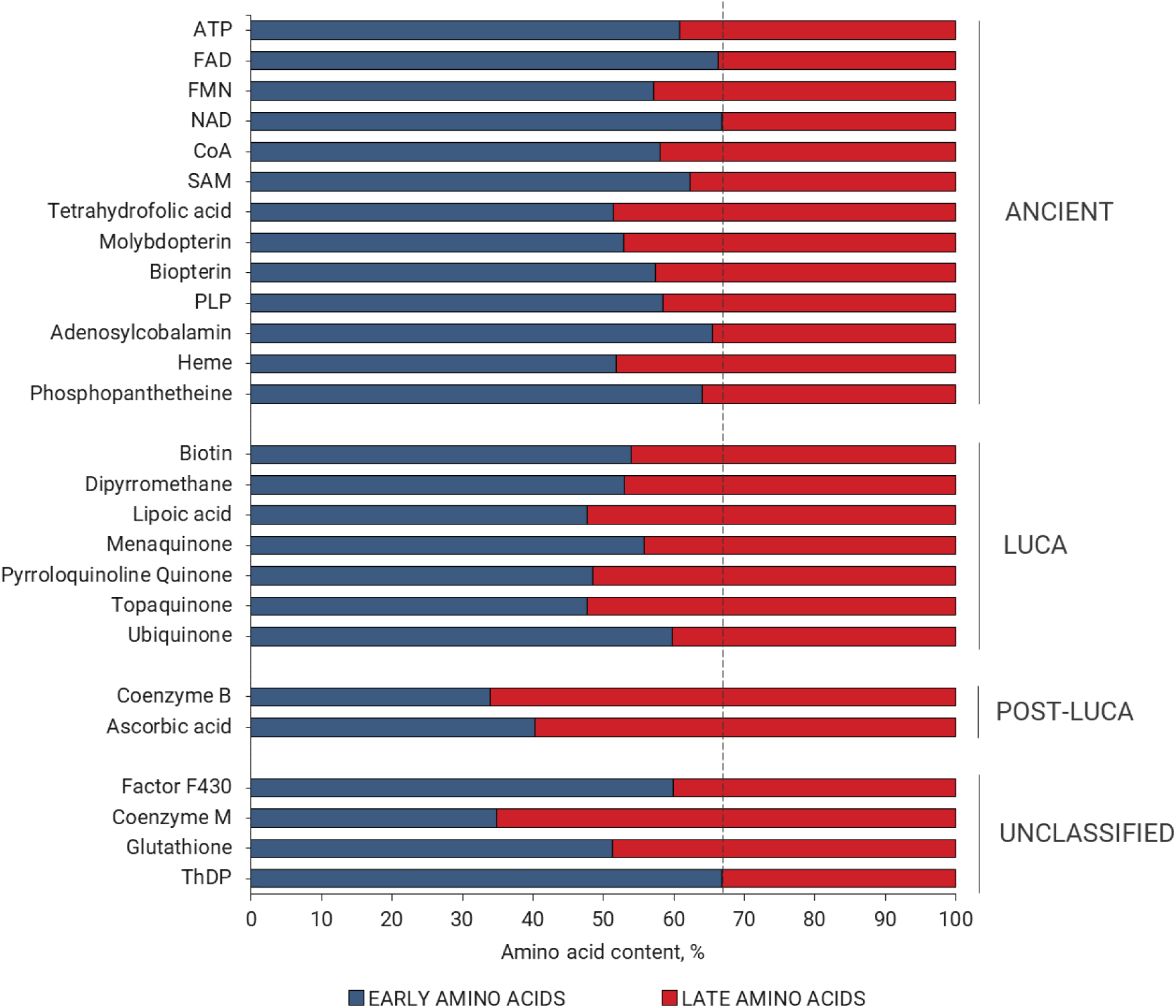
Early versus late amino acid composition of the coenzyme binding sites, categorized according to the evolutionary temporality of coenzymes. Early amino acids are shown in color blue and late residues in red. The dashed line corresponds to the proportion of early vs. late amino acids within the UniProt composition of the sequences derived from our database (67% early and 33% late residues). The statistical significance of the early versus late amino acid composition was assessed by a Chi-squared test (P < 0.0001). Detailed statistical data are listed in Supplementary Table 8.

To explore the contribution of individual amino acids to this effect, fractional difference (FD) for early vs. late amino acids among the Ancient, LUCA, and Post-LUCA coenzyme binding was calculated (Supplementary Table 5A). The mean FD revealed a similar trend to the amino acid composition analysis (Fig. 3). The amino acids most enriched in LUCA vs. Post-LUCA are Gly, Ser, and Leu (FD of 4.4, 4.3, and 4.1 respectively), while the most depleted include Phe, Arg, and His (FD of -11, -4.2, and -3.2) (Supplementary Table 5B).

Moreover, we investigated whether the observed trend in amino acid occurrence at the binding sites was dominated by the presence of phosphate groups, which are common in many Ancient cofactors except for SAM, Tetrahydrofolic acid, Biopterin, and Heme. An additional analysis therefore excluded all phosphate-containing coenzymes indicating that while the trend is less pronounced, it remains even in the absence of phosphate groups (Supplementary Table 6).

To examine the impact that the distribution of amino acids in the coenzyme binding sites has on the binding modes, the interactions between coenzymes and individual amino acids were further inspected.

### Interaction types between coenzymes and proteins

First, backbone vs side chain protein interactions of all the coenzyme classes were mapped (Fig. 4A). As expected, most of the interactions with coenzymes are mediated by amino acid side chains (61 %), frequently in combination with backbone (24%) (Fig. 4). Nevertheless, purely backbone interactions prevail in Ancient coenzymes (24 %) (Fig. 4A). When backbone interactions are present throughout the different coenzyme temporalities, they are dominated by the early amino acids (Fig. 4B).

**Fig 4.**
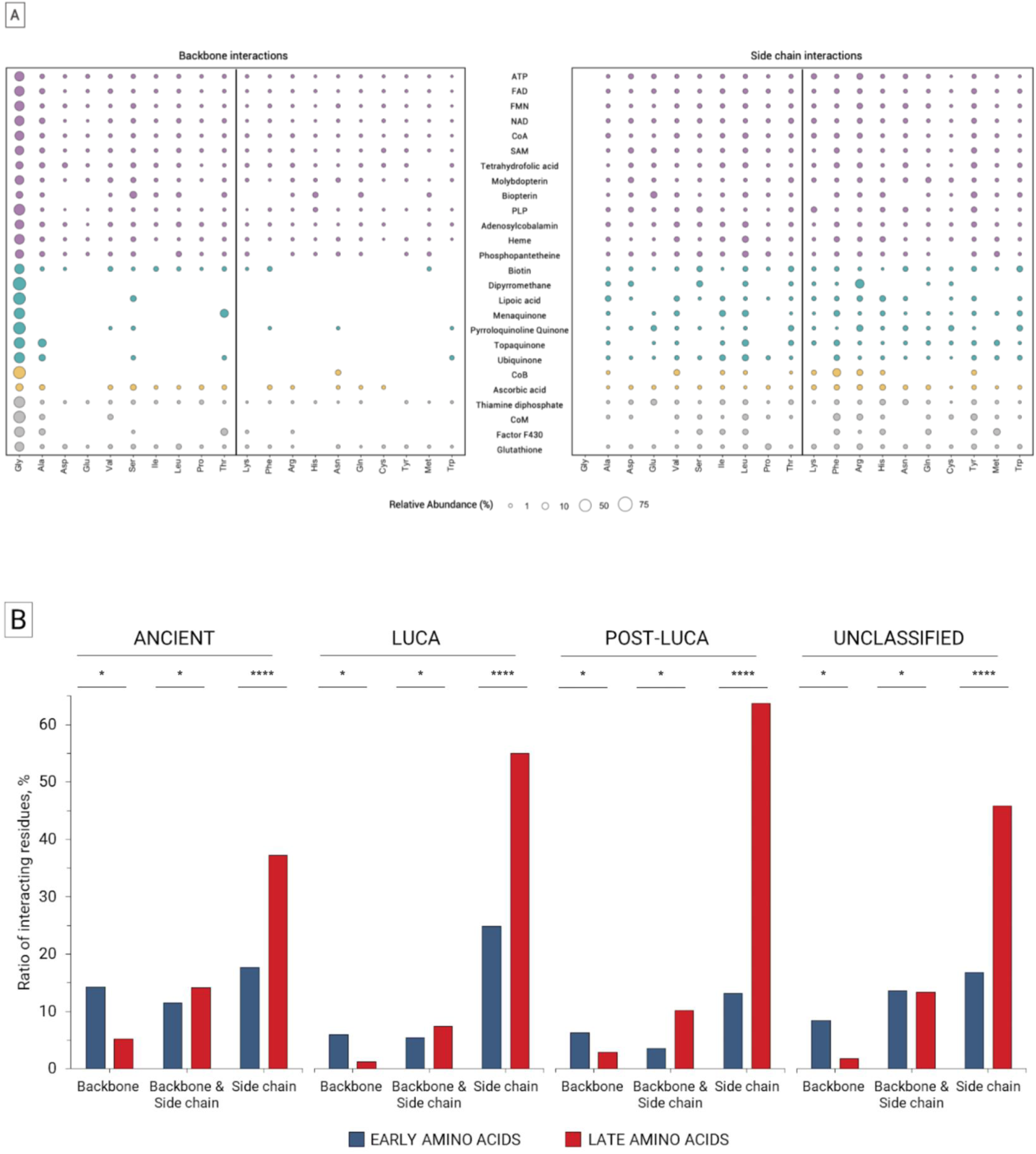
Binding of coenzymes with early and late amino acids by backbone and side chain atoms. "Backbone" interactions refer to residues in the coenzyme binding sites that interact purely through amino acid backbone atoms. "Side chain" interactions involve residues that interact solely via side chain atoms. "Backbone & Side chain" residues are those that interact with the coenzyme using both their backbone and side chain atoms. (A) Abundance of amino acids in individual studied coenzymes. “Backbone & Side chain” interactions are not depicted. Unclassified cofactors are in gray, Post-LUCA in yellow, LUCA in cyan and Ancient in purple. Amino acids are ranked by the order of addition of amino acids to the genetic code (Higgs and Pudritz, 2009). (B) Proportion of early versus late residues in coenzyme categories by interaction type. In each coenzyme category, the individual proportions add up to 100%. The amino acid composition was normalized by the percentage of late residues from the UniProt sequences retrieved from our database. The statistical significance of early versus late amino acid composition for each interaction type per coenzyme temporality was determined by a Chi-squared test (*, P < 0.05; **, P < 0.01; ***, P < 0.001; ****, P < 0.0001) . For detailed statistical analysis, refer to Supplementary Table 9.

Next, we inspected the interaction types for each amino acid-coenzyme binding event employing Arpeggio (Jubb et al., 2017). The analysis revealed that electrostatic interactions are dominant in all coenzyme ages (Supplementary Fig. 2). In Ancient cofactors, electrostatic interactions are more frequently mediated by early residues. This trend is more significant for the nucleotide-derived Ancient coenzymes. The second most prevalent interaction type is Van der Waals for Ancient cofactors while hydrophobic interactions are similarly frequent as Van der Waals in the LUCA and Post-LUCA classes.

### Structural properties of cofactor binding sites

Structural properties of proteins have been observed to change during eons of life’s evolution (Edwards et al., 2013; Lupas and Alva, 2017; Kovacs et al., 2017) To map its interdependence with binding of cofactors, the secondary structure and fold classes of individual coenzyme binding sites were analyzed here.

There are detectable differences in the binding site secondary structure content among the coenzyme temporalities (Fig. 5, Supplementary Fig. 3). While loops and helices dominate all the binding sites, they are less represented in the Ancient and LUCA coenzyme sites which are more rich in beta-sheet structures (Fig. 5). This distinction is found only on the level of the binding sites and not preserved on the level of the overall protein structure.

**Fig 5.**
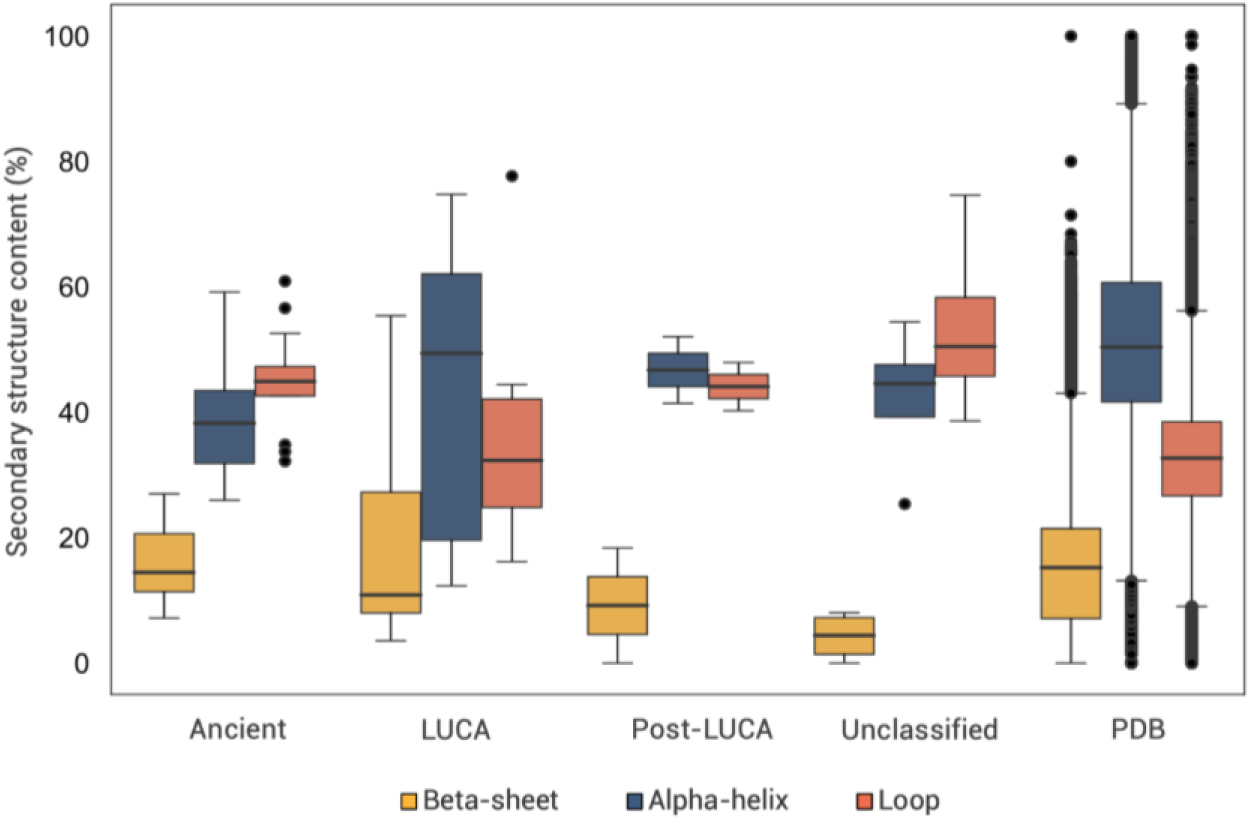
Secondary structure content in coenzyme binding sites. Composition of secondary structural elements in amino acids interacting with coenzymes. The PDB category represents secondary structure content across the dataset for comparison with coenzyme binding sites. Additional statistical analyses are shown in Supplementary Table 10.

To explore the fold diversity of domains containing the coenzyme binding sites, we assigned their ECOD X-groups (Cheng et al., 2014) at a residue level (Supplementary Table 3). In total, 101 groups were identified. The Ancient coenzymes are associated with higher numbers of different X-groups than the LUCA and Post-LUCA cofactors (Fig. 6). Some coenzymes stand out by their large number of associated folds: the Ancient ATP (74); CoA (34); NAD (30); Heme (27) and the Unclassified cofactor Glutathione (25).

**Fig 6.**
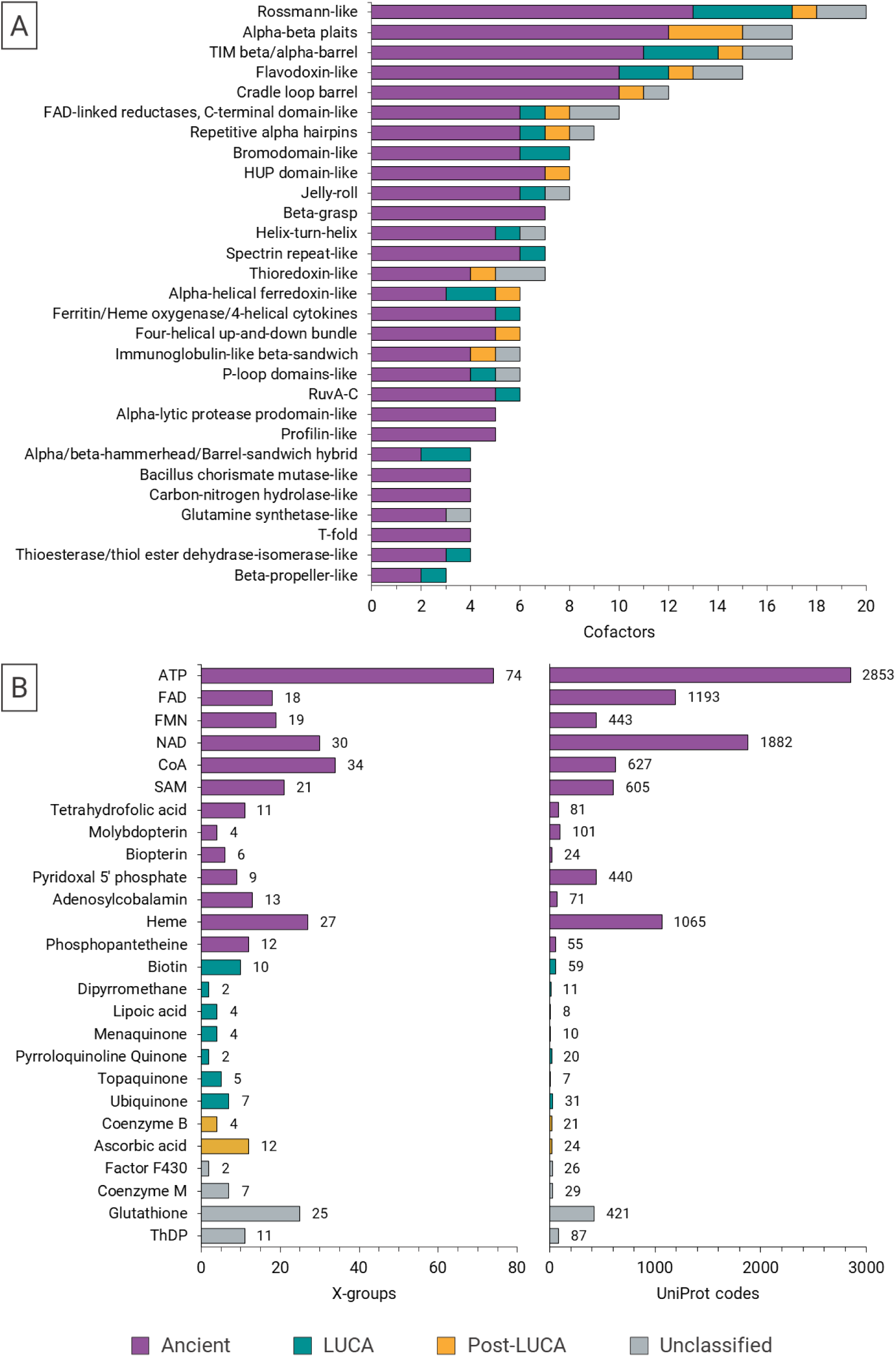
Fold diversity of coenzyme binding sites. (A) Folds represented by ECOD X-groups, according to numbers of coenzyme binding sites. (B) Comparison of numbers of ECOD X-groups vs. UniProt entries per cofactor class

The most frequently observed X-groups include Rossmann-like, Alpha-beta plaits, TIM beta/alpha-barrel, Flavodoxin-like, cradle loop barrel, HUP domain-like and beta-Grasp (Fig. 6). Among these, Rossmann-like, TIM beta/alpha-barrel and Flavodoxin-like bind to coenzymes of all ages.

### Coenzyme early vs late binding sites

To further explore whether extant proteins can bind enzymes only by early or only by late residues (featuring early vs late binding sites), we looked for these specific cases and analyzed their evolutionary conservation.

We found 25 PDB entries that contain at least one chain bound to coenzymes solely by early amino acids (Fig. 7; Supplementary Fig. 4). Those structures correspond to 17 different proteins, represented by unique UniProt codes. The full set of those proteins bind exclusively Ancient coenzymes: ATP, NAD and Phosphopantetheine. In comparison, 15 PDB entries, representing 12 unique proteins, bind coenzymes only by late amino acids (Fig. 7; Supplementary Fig. 4). These examples include all Ancient-to-Post-LUCA coenzymes: ATP, CoA, NAD, PLP, biotin, and Ascorbic acid.

**Fig 7.**
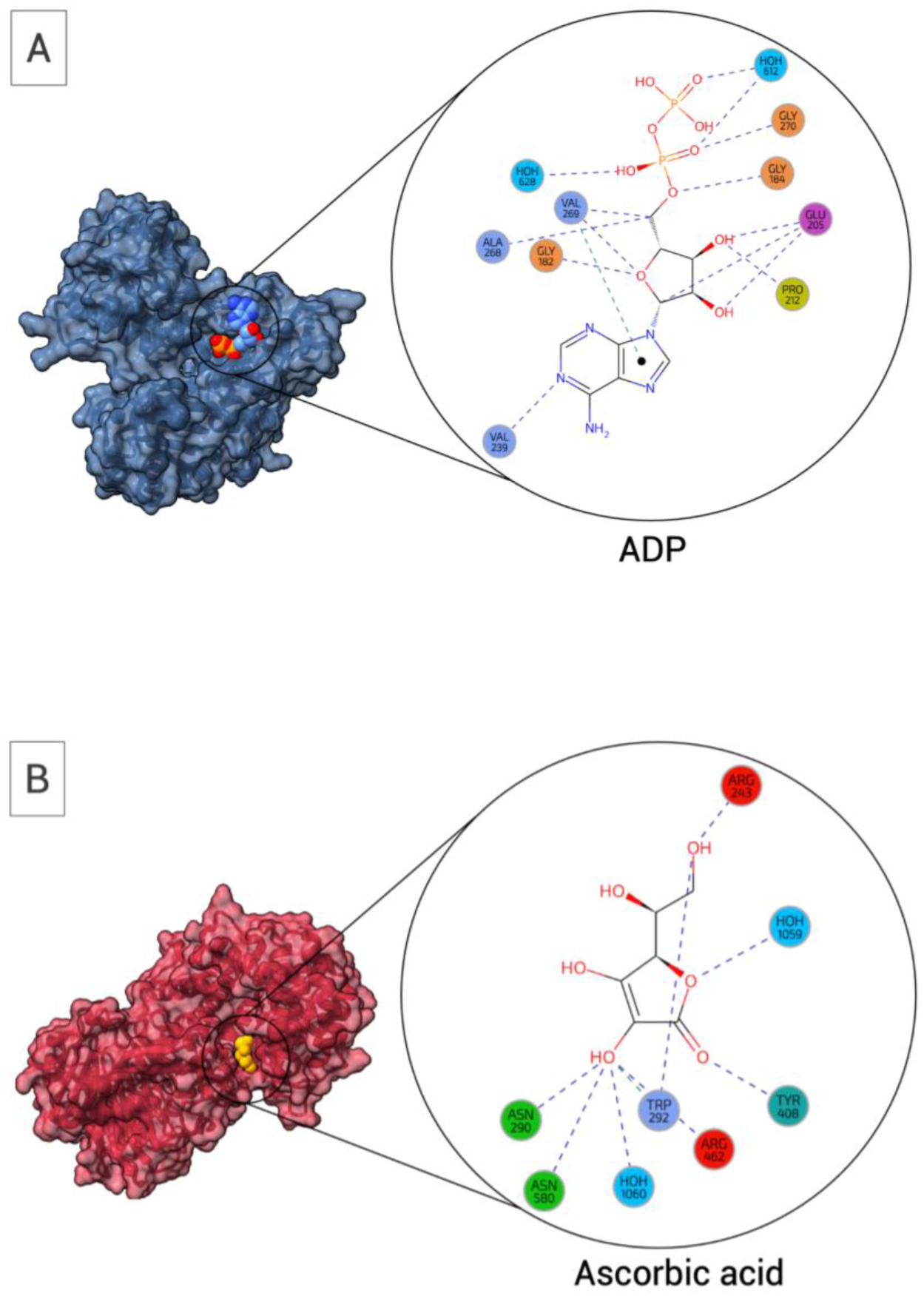
Examples of coenzyme binding solely through early or late amino acids. (A) Coenzymes bound exclusively by early residues (AMP bound by ATP-phosphoribosyltransferase. PDB code 6czm (chain B) created by LIGPLOT (Laskowski and Swindells, 2011). (B) Coenzyme, entirely bound by late residues (Ascorbic acid bound by Hyaluronate lyase. PDB code 1f9g (chain A), created by LIGPLOT).

To assess the conservation of amino acids in these specific binding sites we used ConSurf (Ashkenazy et al., 2010; Ashkenazy et al., 2016). According to the analysis, both the early and late binding sites are relatively highly conserved. Around 60% of the residues from both cases have conservation scores ≥7. Furthermore, we employed the *MAX AA* parameter, that represents the most abundant residue in the multiple sequence alignment of all homologs. 76 vs 72% of the residues in the early vs late binding sites are the same, which suggests their evolutionary conservation in both cases.

### Coenzyme binding mediated by metal ions

Because of the significance of metal ions in both extant and early life, we also analyzed coenzyme binding via metal ions (Supplementary Table 4). Notably, this phenomenon is more frequent in Ancient cofactors, constituting approximately 24% of coenzyme binding sites that have at least one metal ion. Younger cofactors exhibit a lower requirement for metal ion binding. LUCA coenzymes exhibit a metal ion binding in approximately 13% of instances, while in Post-LUCA, 11% and in Unclassified coenzymes, about 23%. Certain coenzymes exhibit a notably high percentage of cases reliant on at least one Mg^2+^ ion (76 % in case of Thiamine diphosphate and 55% in case of ATP binding).

The subsequent most prevalent mediating ion is Ca^2+^, found along with 65 % cases of the LUCA coenzyme Pyrroloquinoline Quinone. Following Ca^2+^, the next most frequent metal ions mediating coenzyme binding are Mn^2+^ and Fe^2+^.

## Discussion

Enzymatic activities rely heavily on interplay with organic cofactors. Those are found at the very heart of cellular metabolism and some of them quite possibly branch deep to life’s early start (White, 1976). The core chemistries of the most abundant and Ancient coenzymes have been repeatedly detected in material and experiments mimicking prebiotic environments (Miller and Schlesinger, 1993; Keefe et al., 1995; Holliday et al., 2007; Kirschning, 2021; Menor-Salván et al., 2022; Pinna et al., 2022; Fairchild et al., 2024). Along with metal ions and minerals, some of the extant coenzymes could probably catalyze metabolic reactions in the absence of enzymes, before their emergence (Muchowska et al. 2020; Henriques Pereira et al. 2022; Cvjetan et al., 2023; Dherbassy et al. 2023). When and how coenzymes seeded the functional hubs of today’s enzymes represents a fundamental bridge between prebiotic chemistry and biochemistry and therefore one of the central questions in the study of life’s origins (Preiner et al. 2020).

The aim of this study was to resolve this conundrum by analyzing protein-coenzyme interactions throughout PDB with respect to amino acid and coenzyme evolutionary temporality (Fig. 2). While no direct inferences about distant evolutionary past can be drawn from the analysis of extant proteins, the principles guiding these interactions can imply their potential prebiotic feasibility and significance.

### Ancient coenzymes are abundant in extant life and bind more frequently to early amino acids

We find that an absolute majority (94 %) of extant coenzymes that appear in PDB structures is conserved across all the life’s domains and available already through prebiotic syntheses (i.e. Ancient). This coenzyme temporality bears many molecules that are derived from nucleotides (such as ATP, FAD, SAM, and NAD) and is spread throughout all the E.C. classes of protein catalysts. This is the first obvious distinction from the other (less-populated) classes of coenzymes and supports the coenzyme-peptide significance from the earliest life until today (White, 1976)

There are several outstanding differences in the properties of binding to proteins among the different coenzyme temporalities. First, the Ancient coenzymes bind to proteins more frequently via early amino acids. On average, this interaction presents 61 % of all Ancient coenzyme bonds to proteins. It is only 53 and 38 % for the LUCA and Post-LUCA coenzymes, respectively.

While the Ancient nucleotide-derived and Unclassified coenzyme binding sites are dominated by residues in the loop conformation (Fig. 5; Supplementary Fig. 3), there is also a substantially higher frequency of residues in beta-sheet conformations when compared to Post-LUCA coenzyme sites. Those, on the other hand, are dominated by alpha-helical conformations. Loops exhibit greater sequence variability compared to ordered structures, and their flexible nature enables them to undergo structural changes (Kessel and Ben-Tal, 2018, Tokuriki and Tawfik, 2009). Such properties of protein active sites were previously associated with evolvability and promiscuity (Tokuriki and Tawfik, 2009, Corbella et al., 2023). The role of loops could thus be important for the flexibility and versatility of early peptide- coenzyme binding sites. It has been noted that evolutionary benefits would be presented by sequences that could adopt closed loop conformations, providing stability and protection to early coenzymes and such hubs could truly transition the peptide-coenzyme world towards primordial enzymes (Goncearenco and Berezovsky 2011, Gamiz-Arco et al., 2021; Toledo-Patiño et al., 2022; Gutierrez-Rus et al., 2023). Besides sequences without regular secondary structure elements, beta-sheets have been considered a more prevalent and significant motif during early stages of protein evolution than alpha helices. Beta-sheet represents the first structural motif in models of the ribosomal evolution, and it has also been observed as a mildly enriched motif in sequences formed from early amino acids (Brack and Orgel 1975, Lupas and Alva 2017, Kovacs et al., 2017, Tretyachenko et al., 2022). Ancient coenzyme-peptide binding properties support the scenario of its significance during early stages of protein evolution.

### Ancient coenzymes bind to proteins through more backbone interactions, typically assigned to early amino acids

While all the coenzymes bind preferentially to protein residue sidechains, more backbone interactions appear in the Ancient coenzyme temporality when compared to others. This supports an earlier hypothesis that functions of the earliest peptides (possibly of variable compositions and lengths) would be performed with the assistance of the main chain atoms rather than their sidechains (Milner-White and Russel 2011). A specific example of such a scenario was recently reported, where a dihydrofolate reductase activity was supported purely by protein backbone-coenzyme interactions (Lemay-St-Denis and Pelletier, 2023). Finally, Longo et al., recently analyzed binding sites of different phosphate-containing ligands which were arguably of high relevance during earliest stages of life, connecting all of today’s core metabolism (Longo et al., 2020 (b)). They observed that unlike the evolutionary younger binding motifs (which rely on sidechain binding), the most ancient lineages indeed bind to phosphate moieties predominantly via the protein backbone.

Our analysis assigns this phenomenon primarily to interactions via early amino acids that (as mentioned above) are generally enriched in the binding interface of the Ancient coenzymes. This implies that late amino acids would not be necessarily needed for the sovereignty of coenzyme-peptide interplay. To address this intriguing possibility, we next searched whether there are such examples in the PDB dataset, where coenzymes would be bound exclusively by early amino acids. We found 17 such proteins where all the coenzymes belonged to the Ancient temporality (such as ATP and NAD). Together with all of the above, this finding supports the possibility that peptide-coenzyme functional hubs could have originated before the evolution of the full canonical amino acid alphabet.

Reinforcing this, we have recently demonstrated on a select example of a ribosomal RNA-binding domain, that a negatively charged variant of the protein composed of only early amino acids is indeed capable of binding to RNA (Giacobelli et al., 2022). In that case, the interaction is further supported by metal ions that were not present at the binding interface of the wild-type protein. Interestingly, the same trend was observed here throughout the PDB dataset. 24% of the Ancient coenzymes in PDB are additionally mediated by at least one metal ion. LUCA and Post-LUCA coenzymes involved metal ions only in 13 and 11 %, respectively. It is quite probable that not all the metal ion densities are recognized or fully resolved in all the PDB structures that were used in our analysis. Nevertheless, we hypothesize that the overall trend can be attributed to the inherent negative charge of many of the Ancient coenzymes, necessitating engagement of positively charged metal ions. This would also be true for direct interaction of early peptides/proteins and metal ions, independent of organic cofactor involvement, as discussed previously by us and others (Bromberg et al., 2022; Frenkel-Pinter et al., 2020; Fried et al., 2022). For example, it has been observed that coordination of prebiotically most relevant metal ions (e.g., Mg^2+^) is more often mediated by early amino acids such as Asp and Glu, whereas metal ions of later relevance (e.g., Cu^+^ and Zn^2+^) bind more frequently via late amino acids like His and Cys (Fried et al. 2022). Similarly, ancient metal binding folds have been shown to be enriched in early amino acids (Bromberg et al., 2022). Along with the general adaptive properties of late amino acids in expanding the chemistry space, the late amino acids would supplement the positive charge in the residue side chains (Ilardo and Freeland, 2014).

### Coenzymes could have served as bridges towards protein structural and functional sovereignty from the peptide-nucleotide world

Our study further revealed that Ancient coenzymes stand out in the variety of protein structures that they bind to, as represented by the ECOD X-groups. While this may partly be caused by their general over-representation in our dataset, the significance of coenzymes has been pointed out previously throughout the most ancient protein folds, such as P-loop NTPases, TIM beta/alpha-barrels, OB and Rossmann folds (Caetano-Anollés et al., 2017; Goldman et al. 2013; Longo et al., 2020(a); Longo et al., 2020(b)). It has been implied by computer simulations that coenzymes could bind to proteins with similar propensity even before the onset of protein homochirality, despite lower structural stability and secondary structure content in heterochiral polypeptides (Skolnick et al., 2019).

Phosphate-containing coenzymes truly stand out by their large number of associated folds (e.g. 74 different ECOD X-groups in case of ATP). It has been postulated that phosphate-binding loops served as the most significant precursors for contemporary enzymes (Romero Romero et al., 2018). Combining ancestral sequence reconstruction and selection/protein design, short polypeptide motifs capable of poly/nucleotide binding have been recovered from the P-loop NTPase and HhH motifs, relying primarily on early amino acids (Longo et al., 2020 (b); Romero Romero et al., 2018). Both demonstrated that these ancient motifs are highly robust to sequence variations, implying that such interaction can be encountered more easily than previously thought (Longo et al., 2020 (b); Keefe and Szostak, 2001). Provokingly, it has also been implied that many of these motifs emerged initially as polynucleotide binders and started serving catalysis only after gaining higher structural complexity (Romero Romero et al., 2018).

Our entry search of PDB coenzymes also retrieved 80 RNA structures. While these were not the primary subject of our analysis, the majority of those were found to interact with Ancient coenzymes, mainly the nucleotide derived ones, such as ATP, FMN, NAD, and SAM (Supplementary Table 7). While some of these structures are assigned as riboswitches, other coenzyme-RNA complexes belong to ribozymes, representing the potential of polynucleotides in early catalysis as discussed by many previous studies (White, 1976; Gilbert, 1986; Reyes-Prieto et al., 2012; Goldman and Kacar 2021). If peptide- polynucleotide interactions were initially more feasible and dominant (in a putative peptide- polynucleotide world, as implied above), coenzymes could have played a key role in resolving the sovereignty of these molecules towards their tertiary structures and catalytic functions. Polypeptide- coenzyme catalysts would soon dominate in performance (enabling more efficient catalysis) and functional repertoires, especially those that would be hard to facilitate in their absence, such as oxidoreductases and transferases (Fried et al., 2022; Goldman and Kacar 2021; Kessel and Ben-Tal 2018).

### Limitations of our work

The first obvious drawback of this analysis is the ambiguity that accompanies the division of coenzymes into evolutionary age temporalities. While the prebiotic availability (or lack of) is quite consensual in some cases, there are also contradictory studies and opinions in the case of some other coenzymes. Some of the coenzymes have e.g. prebiotic precursors but are not present across all the kingdoms of life. This may suggest that the coenzyme became important only Post-LUCA but it can also mean that its importance was only preserved in specific branches of life. “Unclassified” category has been included for such specific cases (presented e.g. by glutathione). Several properties of glutathione that were identified here (such as protein backbone vs. sidechain interaction, ubiquity across all E.C. enzyme classes and a high number of associated X-groups) would suggest that it is closer to Ancient coenzymes. The other “classified” coenzymes were categorized based on prevailing studies although some ambiguity remains. Despite our effort, the evolutionary ratings of coenzymes (and amino acids) are therefore not always clear cut and not all belong to the categories with the same weight (e.g. some are likely to be more ancient than others). For example, the nucleotide-derived coenzymes probably predate the others of the Ancient temporality - it has been proposed that S-adenosylmethionine emerged before the more complex heme-related porphyrin adenosylcobalamin or coenzyme B12 functionalities (Lazcano, 2013).

Another possible bias of our study stems from the population differences among the three coenzyme temporalities. The Ancient coenzymes are by far the most abundant temporality in the PDB dataset. It can be argued that this is the result of their ∼4 billion years of essentiality to life (Goldman and Kacar 2021). Nevertheless, it may as well be contributed by the bias of structures that are deposited in the PDB and most probably does not reflect the true distribution of coenzymes in the biological protein space. Additionally, the comparison of differentially populated temporalities has challenged some aspects of the analysis presented here although care has been taken to perform all the appropriate statistical tests.

### Conclusions

The findings presented here propose that early (the less complex and prebiotically plausible) amino acids are sufficient for binding to Ancient coenzymes. Consequently, coenzyme-peptide interactions might have been conceivable at a time when the amino acid alphabet was not yet evolved to its current form. In such interactions, binding modes would rely more on the protein backbone atoms and on involvement of metal ions, both of which are less frequent in interactions with evolutionarily young coenzymes.

## Methodology

### Identification of organic cofactors

We systematically identified all available cofactor and cofactor-like molecules in the Protein Data Bank in Europe (wwPDB Consortium, 2020) programmatically through the PDBe REST API (pdbe.org/api) (Mukhopadhyay et al., 2019). All cofactor molecules were classified into 27 classes based on the CoFactor database (Fischer et al., 2010). Furthermore, we included ATP and its analogs as an additional cofactor class.

The identification of all the available ligand codes from the PDB chemical component dictionary for each cofactor class was achieved by programmatic access through the “Cofactors” endpoint (https://www.ebi.ac.uk/pdbe/api/pdb/compound/cofactors) using the PDBe REST API (pdbe.org/api) and all responses were in JSON format.

### Structural database and classifications

We retrieved the PDB entries associated to each chemical component from every cofactor class using the “PDB entries containing the compound” endpoint (https://www.ebi.ac.uk/pdbe/api/pdb/compound/in_pdb/:id) via the Entry-based API. The count of PDBe entries for each cofactor class is provided in the supplementary information (Supplementary file 1). The information from the REST API was unavailable for two coenzyme classes, MIO and Orthoquinone, so they were excluded from the analysis.

The secondary structure assignments and (EC) numbers for all PDB structures analyzed were determined through residue-level cross references obtained from the SIFTS XML files (Velankar et al., 2013; Dana et al., 2019). Secondary structure elements include “h” for helix, “b” for strand, and “c” for coil; and they correspond to the information available in the PDBe website. Only observed residues were examined.

Furthermore, we assigned all our PDB entries to the ECOD hierarchical system groups “X”, “H” and “F” (Cheng et al., 2014).

### UniProt assignment and interaction ratio

The assignment of UniProt codes to our structural dataset was achieved by mapping the information with the SIFTS (Velankar et al., 2013; Dana et al., 2019) file “pdb_chain_uniprot.tsv.gz” (https://www.ebi.ac.uk/pdbe/docs/sifts/quick.html). Next, we mapped the UniProt residue to each of the PDB structures with the residue-level cross-reference data of SIFTS by retrieving the XML files (https://www.ebi.ac.uk/pdbe/docs/sifts/quick.html).

Each UniProt code represents a unique protein sequence that encompasses one or various PDBe associated entries. With the aim of filtering those residues relevant to the interaction sites at the level of protein sequence, we incorporated the interaction ratio. The interaction ratio is a measure of the interaction for each ligand with all its PDB associated entries by UniProt residue. Those residue-ligand interactions that were preserved in more than 50% of the associated PDB entries were selected (we call the ratio of preserved interactions among structures of one unique protein interaction ratio).

Upon UniProt residues assignment for each residue of the PDB structures, we downloaded the calculated interaction ratio with the endpoint “UniProt- Get ligand binding residues for a UniProt accession” (https://www.ebi.ac.uk/pdbe/graph-api/uniprot/ligand_sites/:accession).

The redundancy of our database was removed by clustering the UniProt sequences using CD-HIT (Li and Godzik, 2006) with a 90% sequence identity parameter.

### Analysis of coenzyme interactions

In order to analyze the amino acid-coenzyme interactions, we downloaded information of all bound molecules found in a given PDB entry using the “Get bound molecules” endpoint (https://www.ebi.ac.uk/pdbe/graph-api/pdb/bound_molecule_interactions/:pdbId/:bmid) of the Aggregated API. Then, we retrieved the ligand interactions for each bound molecule in every entry through the “PDB-Get bound ligand interactions” endpoint (https://www.ebi.ac.uk/pdbe/graph-api/pdb/bound_ligand_interactions/:pdbId/:chain/:seqId), which calculates these interactions with Arpeggio (Jubb et al., 2017). The retrieved interactions included the standard amino acid codes, water molecules and metal ions.

We classified all the interactions reported by Arpeggio into nine distinct interaction types. The classification scheme aligns with the one used by PDBe and encompasses the following categories: i) “covalent”; ii) "electrostatic", which combines "ionic", "hbond", "weak_hbond", "polar", "weak_polar", "xbond" and "carbonyl"; iii) “amide”, consisting in "AMIDEAMIDE" and "AMIDERING"; iv) "vdw", denoting van der Waals interactions; v) "hydrophobic"; vi) "aromatic", grouping "aromatic", "FF", "OF", "EE", "FT", "OT", "ET", "FE", "OE" and "EF" contacts; vii) "atom-pi", comprised of "CARBONPI", "CATIONPI", "DONORPI", "HALOGENPI", and "METSULPHURPI"; viii) "metal" and ix) "clashes", including "clash" and "vdw_clash" contacts. We have omitted this last category due to the limited number of interactions, most of which result from experimental errors during X-ray diffraction.

Backbone and side chains interactions were identified based on the atom identities in the coenzyme binding sites. Those atoms corresponding to the backbone of standard amino acids were identified as: "N”, “C", "CA", "O". Glycine has only a hydrogen atom as its side chain; nevertheless, no side chain atom mediating any interaction was identified.

### Secondary structure analysis

Statistical analysis of secondary structure content was conducted at the UniProt level. For each residue within every UniProt entry, we considered all potential secondary structure elements derived from the PDB structures associated with each UniProt code. Subsequently, we eliminated redundancy on a per-residue basis. This methodological approach enabled us to comprehensively encompass the structural diversity at each position of the protein.

### Interactions mediated exclusively by early or late amino acids

To examine proteins that interacted with cofactors solely through early or late amino acids, we filtered the data to include only proteins interacting with at least two amino acids.

For the assessment of the evolutionary conservation of coenzyme-binding amino acids, we employed ConSurf (Ashkenazy et al., 2010; Ashkenazy et al., 2016). Specifically, we analyzed the msa_positional_aa_frequency files generated for each PDB structure.

## Acknowledgements

This work was supported by the Human Frontier Science Program grant HFSP-RGEC27/2023 and was carried out with the support of ELIXIR CZ Research Infrastructure (ID LM2023055, MEYS CR). A.C.S.R. and M.M. acknowledge support by the project the “Grant Schemes at CU” (reg. no. CZ.02.2.69/0.0/0.0/19_073/0016935), project no. START/SCI/148. Finally, we would like to thank Prof. Stephen Freeland and Prof. Janet Thornton for helpful discussions on this manuscript.

## Notes

### Competing Interest Statement

The authors have declared no competing interest.

### Summary of Updates

Results have been extended by current Suppl Tables 5&6 and discussion has been extended

## References

1. Alva V, Söding J, Lupas AN. (2015). A vocabulary of ancient peptides at the origin of folded proteins. eLife 4:e09410. 10.7554/eLife.09410, PMID: 26653858

2. Aylward N. (2006). An ab initio computational study of thiamin synthesis from gaseous reactants of the interstellar medium. Biophysical Chemistry 121: 185–193. DOI:10.1016/j.bpc.2005.12.018, PMID: 16527390

3. Aylward, N, Bofinger N. (2006). A plausible prebiotic synthesis of pyrdoxal phosphate: Vitamin B-6 - A computational study. Biophysical Chemistry 123: 113–121. DOI:10.1016/j.bpc.2006.04.014, PMID: 16730878

4. Ashkenazy H, Erez E, Martz E, Pupko T, Ben-Tal N. (2010). ConSurf 2010: calculating evolutionary conservation in sequence and structure of proteins and nucleic acids. Nucleic Acids Res 38:W529–33. DOI:10.1093/nar/gkq399, PMID: 20478830

5. Ashkenazy H, Abadi S, Martz E, Chay O, Mayrose I, Pupko T, Ben-Tal N. (2016). ConSurf 2016: an improved methodology to estimate and visualize evolutionary conservation in macromolecules. Nucleic Acids Res 44(W1):W344–50. DOI:10.1093/nar/gkw408, PMID: 27166375

6. Barrick JE, Breaker R. (2007). The distributions, mechanisms, and structures of metabolite-binding riboswitches. Genome Biology 8(11):R239. 10.1186/gb-2007-8-11-r239, PMID: 17997835

7. Bonfio C, Valer L, Scintilla S, Shah S, Evans D, Jin L, Szostak J, Sasselov D, Sutherland J, Mansy S. (2017). UV-light-driven prebiotic synthesis of iron-sulfur clusters. Nature Chemistry, 9(12): 1229– 1234. 10.1038/nchem.2817, PMID: 29168482

8. Burton AS, Stern JC, Elsila JE, Glavin DP, Dworkin JP. (2012). Understanding prebiotic chemistry through the analysis of extraterrestrial amino acids and nucleobases in meteorites. Chemical Society reviews 41(16): 5459–72. DOI:10.1039/c2cs35109a, PMID: 22706603

9. Brack A, Orgel LE. (1975). Beta structures of alternating polypeptides and their possible prebiotic significance. Nature 256(5516):383–7. DOI:10.1038/256383a0, PMID: 238134

10. Bromberg Y, Aptekmann AA, Mahlich Y, Cook L, Senn S, Miller M, Nanda V, Ferreiro DU, Falkowski PG. (2022). Quantifying structural relationships of metal-binding sites suggests origins of biological electron transfer. Science Advances. 8(2). 10.1126/sciadv.abj3984

11. Caetano-Anollés G., Hee SK, Mittenthal JE. (2007). The origin of modern metabolic networks inferred from phylogenomic analysis of protein architecture. Proceedings of the National Academy of Sciences of the United States of America 104(22):9358–9363. 10.1073/pnas.0701214104, PMID: 17517598

12. Cheng H, Schaeffer RD, Liao Y, Kinch LN, Pei J, Shi S, Kim BH, Grishin NV. (2014). ECOD: an evolutionary classification of protein domains. PLoS Computational Biol 10(12):e1003926. DOI:10.1371/journal.pcbi.1003926. PMID: 25474468

13. Chu XY, Zhang HY. (2020). Cofactors as molecular fossils to trace the origin and evolution of proteins. ChemBioChem **21**:3161–3168. DOI:10.1002/cbic.202000027, PMID: 32515532

14. Cvjetan N, Schuler L, Ishikawa T, Walde P. (2023). Optimizationand Enhancementof the Peroxidase- like Activity of Hemin in Aqueous Solutions of Sodium Dodecylsulfate. ACS Omega 8(45):42878–42899. DOI: 10.1021/acsomega.3c05915

15. Cleaves HJ. (2010). The origin of the biologically coded amino acids. Journal of Theoretical Biology 263(4): 490–498. 10.1016/j.jtbi.2009.12.014, PMID: 20034500

16. Copley SD, Dhillon JK. (2002). Lateral gene transfer and parallel evolution in the history of glutathione biosynthesis genes. Genome Biology 3(5): 1–16. 10.1186/gb-2002-3-5-research0025, PMID: 12049666

17. Corbella M, Pinto GP, Kamerlin SCL. (2023). Loop dynamics and the evolution of enzyme activity. Nature Reviews Chemistry 7(8):536–547. 10.1038/s41570-023-00495-w, PMID: 37225920

18. Dana JM, Gutmanas A, Tyagi N, Qi G, O’Donovan C, Martin M, Velankar S. (2019). SIFTS: updated Structure Integration with Function, Taxonomy and Sequences resource allows 40-fold increase in coverage of structure-based annotations for proteins. Nucleic Acids Res **47(D1):**D482-D489. DOI:10.1093/nar/gky1114, PMID: 30445541

19. Dherbassy Q, Mayer R, Muchowska K, & Moran J. (2023). Metal-Pyridoxal Cooperativity in Nonenzymatic Transamination. Journal of the American Chemical Society 145 (24):13357–13370. DOI: 10.1021/jacs.3c03542, PMID: 37278531

20. Edwards H, Abeln S, Deane CM. (2013). Exploring Fold Space Preferences of New-born and Ancient Protein Superfamilies. PLoS Computational Biology 9(11):e1003325. 10.1371/journal.pcbi.1003325, PMID: 24244135

21. Fairchild J, Islam S, Singh J, Powner MW. (2024). Prebiotically plausible chemoselective pantetheine synthesis in water. Science 383:911–918. DOI: 10.1126/science.adk4432

22. Fischer JD, Holliday GL, Thornton JM. (2010). The CoFactor database: Organic cofactors in enzyme catalysis. Bioinformatics 26(19):2496–2497. 10.1093/bioinformatics/btq442, PMID: 20679331

23. Frenkel-Pinter M, Haynes JW, Martin C, Petrov AS, Burcar BT, Krishnamurthy R, Hud N, Leman L, Williams LD. (2019). Selective incorporation of proteinaceous over nonproteinaceous cationic amino acids in model prebiotic oligomerization reactions. Proceedings of the National Academy of Sciences of the United States of America 116(33):16338–16346. 10.1073/pnas.1904849116, PMID: 31358633

24. Frenkel-Pinter M, Mousumi S, Ashkenasy G, Leman L. (2020). Prebiotic Peptides: Molecular Hubs in the Origin of Life. Chemical Reviews 120(11): 4707–4765. DOI: 10.1021/acs.chemrev.9b00664, PMID: 32101414

25. Fried SD, Fujishima K, Makarov M, Cherepashuk I, Hlouchova K. (2022). Peptides before and during the nucleotide world: An origins story emphasizing cooperation between proteins and nucleic acids. Journal of the Royal Society Interface 19(187):20210641. DOI: 10.1098/rsif.2021.0641, PMID: 35135297

26. Gamiz-Arco G, Gutierrez-Rus LI, Risso VA, Ibarra-Molero B, Hoshino Y, Petrović D, Justicia J, Cuerva JM, Romero-Rivera A, Seelig B, Gavira JA, Kamerlin SCL, Gaucher EA, Sanchez-Ruiz JM. (2021). Heme-binding enables allosteric modulation in an ancient TIM-barrel glycosidase. Nature Communications 12(1):380. 10.1038/s41467-020-20630-1, PMID: 33452262

27. Giacobelli, VG, Fujishima K, Lepšík M, Tretyachenko V, Kadavá T, Makarov M, Hlouchová, K. (2022). In Vitro Evolution Reveals Noncationic Protein-RNA Interaction Mediated by Metal Ions. Molecular Biology and Evolution 39(3):1–11. 10.1093/molbev/msac032, PMID: 35137196

28. Gilbert W. (1986). The RNA world superlattices point ahead. Nature 319: 618.

29. Goldman AD, Bernhard TM, Dolzhenko E, Landweber LF. (2013). LUCApedia: A database for the study of ancient life. Nucleic Acids Res 41(D1): 1079–1082. 10.1093/nar/gks1217, PMID: 23193296

30. Goldman AD, Kacar B. (2021). Cofactors are Remnants of Life’s Origin and Early Evolution. Journal of Molecular Evolution 89(3):127–133. DOI: 10.1007/s00239-020-09988-4, PMID: 33547911

31. Goncearenco A, Berezovsky IN. (2011). Prototypes of elementary functional loops unravel evolutionary connections between protein functions. Bioinformatics 27(13):i497–i503. 10.1093/bioinformatics/btq374, PMID: 20823313

32. Gutierrez-Rus LI, Gamiz-Arco G, Gavira JA, Gaucher EA, Risso VA, Sanchez-Ruiz JM. (2023). Protection of Catalytic Cofactors by Polypeptides as a Driver for the Emergence of Primordial Enzymes. Molecular Biology and Evolution 40(6):1–8. 10.1093/molbev/msad126, PMID: 37235753

33. Henriques DP, Leethaus J, Beyazay T, do Nascimento A, Kleinermanns K, Tüysüz H, Martin W, Preiner M. (2022). Role of geochemical protoenzymes (geozymes) in primordial metabolism: specific abiotic hydride transfer by metals to the biological redox cofactor NAD+. FEBS Journal **289(11)**:3148–3162. 10.1111/febs.16329, PMID: 34923745

34. Higgs PG, Pudritz RE. (2009). A thermodynamic basis for prebiotic amino acid synthesis and the nature of the first genetic code. Astrobiology 9(5):483–490. 10.1089/ast.2008.0280, PMID: 19566427

35. Holliday GL, Thornton JM, Marquet A, Smith AG, Rébeillé F, Mendel R, Schubert HL, Lawrence AD, Warren MJ. (2007). Evolution of enzymes and pathways for the biosynthesis of cofactors. Natural Product Reports 24(5): 972–987. 10.1039/b703107f, PMID: 17898893

36. Huang F, Bugg CW, Yarus M. RNA-Catalyzed CoA, NAD, and FAD synthesis from phosphopantetheine, NMN, and FMN. (2000). Biochemistry **39(50)**:15548-55. DOI: 10.1021/bi002061f, PMID: 11112541

37. Ilardo MA, Freeland SJ. (2014). Testing for adaptive signatures of amino acid alphabet evolution using chemistry space. Journal of Systems Chemistry 5(1):1–9. 10.1186/1759-2208-5-1

38. Ji HF, Chen L, Zhang HY. (2008). Organic cofactors participated more frequently than transition metals in redox reactions of primitive proteins. BioEssays 30(8):766–771. 10.1002/bies.20788, PMID: 18618622

39. Jubb HC, Higueruelo AP, Ochoa-Montaño B, Pitt WR, Ascher DB, Blundell TL. (2017). Arpeggio: A Web Server for Calculating and Visualising Interatomic Interactions in Protein Structures. J Mol Biol **429(3)**:365-371. DOI: 10.1016/j.jmb.2016.12.004, PMID: 27964945

40. Keefe AD, Newton GL, Miller SL. (1995). A possible prebiotic synthesis of pantetheine, a precursor to coenzyme a. Nature 373(6516): 683–685. 10.1038/373683a0, PMID: 7854449

41. Keefe AD, Szostak JW. (2001). Functional proteins from a random-sequence library. Nature 410(6829):715–8. DOI: 10.1038/35070613. PMID: 11287961

42. Kessel A, Ben-Tal N. (2018). Introduction to proteins: structure, function, and motion. Crc Press (Taylor & Francis Group). DOI:10.1201/9781315113876

43. Kessel A, Ben-Tal N. (2022). From Molecules to Cells: The Origin of Life on Earth. Kindle E-Book

44. Kirschning A. (2021). Coenzymes and Their Role in the Evolution of Life. Angewandte Chemie - International Edition 60(12):6242–6269. 10.1002/anie.201914786, PMID: 31945250

45. Kirschning A. (2022). On the Evolutionary History of the Twenty Encoded Amino Acids. Chemistry - A European Journal 28(55):e202201419, 10.1002/chem.202201419, PMID: 35726786

46. Kolodny R, Nepomnyachiy S, Tawfik DS, Ben-Tal N. (2021). Bridging Themes: Short Protein Segments Found in Different Architectures. Molecular Biology and Evolution 38(6): 2191–2208. 10.1093/molbev/msab017, PMID: 33502503

47. Kovacs NA, Petrov AS, Lanier KA, Williams LD. (2017). Frozen in Time: The History of Proteins. Molecular Biology and Evolution 34(5):1252–1260. 10.1093/molbev/msx086, PMID: 28201543

48. Lane N, Martin WF. (2012). The origin of membrane bioenergetics. Cell 151(7):1406–1416. 10.1016/j.cell.2012.11.050, PMID: 23260134

49. Laurino P, Tóth-Petróczy Á, Meana-Pañeda, Lin W, Truhlar DG, Tawfik DS. (2016). An Ancient Fingerprint Indicates the Common Ancestry of Rossmann-Fold Enzymes Utilizing Different Ribose-Based Cofactors. PLoS Biology 14(3): 1–23. 10.1371/journal.pbio.1002396, PMID: 26938925

50. Laskowski RA, Swindells MB. (2011). LigPlot+: multiple ligand-protein interaction diagrams for drug discovery. J Chem Inf Model 51(10):2778–86. DOI:10.1021/ci200227u, PMID: 21919503

51. Lazcano A. (2013). Planetary change and biochemical adaptation: Molecular evolution of corrinoid and heme biosyntheses. Hematology 17:s7–s10. 10.1179/102453312X13336169155015

52. Lemay-St-Denis C, Pelletier J. (2023). From a binding module to essential catalytic activity: how nature stumbled on a good thing. Chem. Commun 59:12560–12572. 10.1039/D3CC04209J

53. Li W, Godzik A. (2006). Cd-hit: a fast program for clustering and comparing large sets of protein or nucleotide sequences. Bioinformatics 22(13):1658–9. DOI:10.1093/bioinformatics/btl158, PMID: 16731699

a. Longo LM, Jabtoiiska J, Vyas P, Kanade M, Kolodny R, Ben-Tal N, Tawfik DS. (2020). On the emergence of p-loop ntpase and rossmann enzymes from a beta-alpha-beta ancestral fragment. Elife 9:1–16. 10.7554/ELIFE.64415, PMID: 33295875

b. Longo LM, Petrovic D, Kamerlin SCL, Tawfik DS. (2020). Short and simple sequences favored the emergence of N-helix phospho-ligand binding sites in the first enzymes. Proceedings of the National Academy of Sciences of the United States of America 117(10):5310–5318. 10.1073/pnas.1911742117, PMID: 32079722

56. Lupas AN, Alva V. (2017). Ribosomal proteins as documents of the transition from unstructured (poly)peptides to folded proteins. Journal of Structural Biology 198(2):74–81. 10.1016/j.jsb.2017.04.007, PMID: 28454764

57. Makarov M, Meng J, Tretyachenko V, Srb P, Březinová A, Giacobelli VG, Bednárová L, Vondrášek J, Dunker K, Hlouchová K. (2021). Enzyme catalysis prior to aromatic residues: Reverse engineering of a dephospho-CoA kinase. Protein Science 30(5):1022–1034. 10.1002/pro.4068, PMID: 33739538

58. Menor-Salván C, Burcar BT, Bouza M, Fialho DM, Fernández FM, Hud NV. (2022). A Shared Prebiotic Formation of Neopterins and Guanine Nucleosides from Pyrimidine Bases. Chemistry (Weinheim an Der Bergstrasse, Germany) 28(39):e202200714. 10.1002/chem.202200714, PMID: 35537135

59. Miller SL, Schlesinger G. (1993). Prebiotic syntheses of vitamin coenzymes: I. Cysteamine and 2- mercaptoethanesulfonic acid (coenzyme M). Journal of Molecular Evolution **36(4):**302-7. DOI: 10.1007/BF00182177, PMID: 11536534

60. Milner-White EJ, Russell MJ. Functional capabilities of the earliest peptides and the emergence of life. (2011). Genes. **2(4):**671-88. DOI:10.3390/genes2040671. PMID: 24710286

61. Monteverde DR, Gómez-Consarnau L, Suffridge C, Sañudo-Wilhelmy SA. (2017). Life’s utilization of B vitamins on early Earth. Geobiology 15(1):3–18. 10.1111/gbi.12202

62. Muchowska KB, Varma SJ, Moran J. (2020). Nonenzymatic Metabolic Reactions and Life’s Origins. Chemical Reviews, 120(15):7708–7744. 10.1021/acs.chemrev.0c00191, PMID: 32687326

63. Mukhopadhyay A, Borkakoti N, Pravda L, Tyzack JD, Thornton JM, Velankar S. (2019). Finding enzyme cofactors in Protein Data Bank. Bioinformatics 35(18):3510–3511. 10.1093/bioinformatics/btz115, PMID: 32687326

64. Naraoka H, Takano Y, Dworkin JP, Oba Y, Hamase K, Furusho A, Ogawa NO, Hashiguchi M, Fukushima K, Aoki D, Schmitt-Kopplin P, Aponte JC, Parker ET, Glavin DP, McLain HL, Elsila JE, Graham HV, Eiler JM, Orthous-Daunay FR, Wolters C, Isa J, Vuitton V, Thissen R, Sakai S, Yoshimura T, Koga T, Ohkouchi N, Chikaraishi Y, Sugahara H, Mita H, Furukawa Y, Hertkorn N, Ruf A, Yurimoto H, Nakamura T, Noguchi T, Okazaki R, Yabuta H, Sakamoto K, Tachibana S, Connolly HC Jr, Lauretta DS, Abe M, Yada T, Nishimura M, Yogata K, Nakato A, Yoshitake M, Suzuki A, Miyazaki A, Furuya S, Hatakeda K, Soejima H, Hitomi Y, Kumagai K, Usui T, Hayashi T, Yamamoto D, Fukai R, Kitazato K, Sugita S, Namiki N, Arakawa M, Ikeda H, Ishiguro M, Hirata N, Wada K, Ishihara Y, Noguchi R, Morota T, Sakatani N, Matsumoto K, Senshu H, Honda R, Tatsumi E, Yokota Y, Honda C, Michikami T, Matsuoka M, Miura A, Noda H, Yamada T, Yoshihara K, Kawahara K, Ozaki M, Iijima YI, Yano H, Hayakawa M, Iwata T, Tsukizaki R, Sawada H, Hosoda S, Ogawa K, Okamoto C, Hirata N, Shirai K, Shimaki Y, Yamada M, Okada T, Yamamoto Y, Takeuchi H, Fujii A, Takei Y, Yoshikawa K, Mimasu Y, Ono G, Ogawa N, Kikuchi S, Nakazawa S, Terui F, Tanaka S, Saiki T, Yoshikawa M, Watanabe SI, Tsuda Y. (2023). Soluble organic molecules in samples of the carbonaceous asteroid (162173) Ryugu. Science 379(6634):eabn9033. 10.1126/science.abn9033, PMID: 36821691

65. Narunsky A, Kessel A, Solan R, Alva V, Kolodny R, Ben-Tal N. (2020). On the evolution of protein- adenine binding. Proceedings of the National Academy of Sciences of the United States of America 117(9):4701–4709. 10.1073/pnas.1911349117, PMID: 32079721

66. Nepomnyachiy S, Ben-Tal N, Kolodny R. (2017). Complex evolutionary footprints revealed in an analysis of reused protein segments of diverse lengths. Proceedings of the National Academy of Sciences of the United States of America. 114(44):11703–11708. DOI: 10.1073/pnas.1707642114, PMID: 29078314

67. PDBe-KB consortium. (2020). PDBe-KB: a community-driven resource for structural and functional annotations. Nucleic Acids Res 48(D1):D344–D353. DOI:10.1093/nar/gkz853, PMID: 31584092

68. Pinna S, Kunz C, Halpern A, Harrison SA, Jordan SF, Ward J, Werner F, Lane N. (2022). A prebiotic basis for ATP as the universal energy currency. PLoS Biology 20(10):1–25. 10.1371/journal.pbio.3001437, PMID: 36194581

69. Putignano V, Rosato A, Banci L, Andreini C. (2018). MetalPDB in 2018: A database of metal sites in biological macromolecular structures. Nucleic Acids Res 46(D1):D459–D464. 10.1093/nar/gkx989, PMID: 29077942

70. Preiner M, Asche S, Becker S, Betts HC, Boniface A, Camprubi E, Chandru K, Erastova V, Garg SG, Khawaja N, Kostyrka G, Machné R, Moggioli G, Muchowska KB, Neukirchen S, Peter B, Pichlhöfer E, Radványi Á, Rossetto D, Salditt A, Schmelling NM, Sousa FL, Tria FDK, Vörös D, Xavier JC. (2020). The future of origin of life research: Bridging decades-old divisions. Life 10(3): 20. 10.3390/life10030020, PMID: 32110893

71. Qiu K, Ben-Tal N, Kolodny R. (2022). Similar protein segments shared between domains of different evolutionary lineages. Protein Science 31(9):e4407. DOI:10.1002/pro.4407. PMID: 36040261

72. Reyes-Prieto F, Hernández-Morales R, Jácome R, Becerra A, Lazcano A. (2012). Coenzymes, viruses and the RNA world. Biochimie 94(7):1467–1473. 10.1016/j.biochi.2012.01.004, PMID: 22269935

73. Romero Romero ML, Yang F, Lin YR, Toth-Petroczy A, Berezovsky IN, Goncearenco A, Yang W, Wellner A, Kumar-Deshmukh F, Sharon M, Baker D, Varani G, Tawfik DS. (2018). Simple yet functional phosphate-loop proteins. Proceedings of the National Academy of Sciences of the United States of America 115(51):E11943–E11950. 10.1073/pnas.1812400115, PMID: 30504143

74. Russell MJ, Hall AJ. (1997). The emergence of life from iron monosulphide bubbles at a submarine hydrothermal redox and pH front. J Geol Soc London 154(3):377–402. DOI: 10.1144/gsjgs.154.3.0377. PMID:11541234

75. Seitz C, Eisenreich W, Huber C. (2021). The abiotic formation of pyrrole under volcanic, hydrothermal conditions—an initial step towards life’s first breath? Life 11(9):1–10. 10.3390/life11090980, PMID: 34575129

76. Skolnick J, Zhou H, Gao M. (2019). On the possible origin of protein homochirality, structure, and biochemical function. Proceedings of the National Academy of Sciences of the United States of America 116(52): 26571–26579. 10.1073/pnas.1908241116

77. Söding J, Lupas AN. (2003). More than the sum of their parts: On the evolution of proteins from peptides. BioEssays 25(9):837–846. 10.1002/bies.10321, PMID: 12938173

78. Thauer RK, Bonacker LG. (1994). Biosynthesis of coenzyme F430, a nickel porphinoid involved in methanogenesis. Ciba Found Symp 180:210–22. DOI: 10.1002/9780470514535.ch12. PMID: 7842854

79. The UniProt Consortium. (2023) UniProt: the Universal Protein Knowledgebase in 2023. Nucleic Acids Res 51:D523–D531. 10.1093/nar/gkac1052, PMID: 36408920

80. Toledo-Patiño S, Pascarelli S, Uechi GI, Laurino P. (2022). Insertions and deletions mediated functional divergence of Rossmann fold enzymes. Proceedings of the National Academy of Sciences of the United States of America 119(48):e2207965119. DOI: 10.1073/pnas.2207965119, PMID: 36417431

81. Tokuriki N, Tawfik DS. (2009). Protein dynamism and evolvability. Science 324(5924):203–7. DOI: 10.1126/science.1169375. PMID: 19359577

82. Tretyachenko V, Vymětal J, Neuwirthová T, Vondrášek J, Fujishima K, Hlouchová K. (2022). Modern and prebiotic amino acids support distinct structural profiles in proteins. Open Biol 12(6):220040. DOI: 10.1098/rsob.220040, PMID: 35728622

83. Trifonov EN. (2000). Consensus temporal order of amino acids and evolution of the triplet code. Gene 261(1):139–151. 10.1016/S0378-1119(00)00476-5, PMID: 11164045

84. Velankar S, Dana JM, Jacobsen J, van Ginkel G, Gane PJ, Luo J, Oldfield TJ, O’Donovan C, Martin MJ, Kleywegt GJ. (2013). SIFTS: Structure Integration with Function, Taxonomy and Sequences resource. Nucleic Acids Res 41:D483–9. DOI: 10.1093/nar/gks1258, PMID: 23203869

85. Wächtershäuser G. (1992). Groundworks for an evolutionary biochemistry: The iron-sulphur world. Progress in Biophysics and Molecular Biology 58(2):85–201. 10.1016/0079-6107(92)90022-X, PMID: 1509092

86. Weber AL, Miller SL. (1981). Reasons for the occurrence of the twenty coded protein amino acids. Journal of Molecular Evolution 17(5):273–84. DOI:10.1007/BF01795749, PMID: 7277510

87. White HB. (1976). Coenzymes as fossils of an earlier metabolic state. Journal of Molecular Evolution 7(2):101–104. 10.1007/BF01732468, PMID: 1263263

88. White HB. (1982). Evolution of Coenzymes and the Origin of Pyridine Nucleotides. The Pyridine Nucleotide Coenzymes. Econometrica 50:1–17. 10.1016/b978-0-12-244750-1.50010-5

89. Wong JT, Bronskill PM. (1979). Inadequacy of prebiotic synthesis as origin of proteinous amino acids. Journal of Molecular Evolution 13(2):115–25. DOI:10.1007/BF01732867, PMID: 480369

90. Wu HH, Pun MD, Wise CE, Streit BR, Mus F, Berim A, Kincannon WM, Islam A, Partovi SE, Gang DR, DuBois JL, Lubner CE, Berkman CE, Lange BM, Peters JW. (2022). The pathway for coenzyme M biosynthesis in bacteria. Proceedings of the National Academy of Sciences of the United States of America 119(36):e2207190119. DOI:10.1073/pnas.220719011, PMID: 36037354

91. Zaia DA, Zaia CT, De Santana H. Which amino acids should be used in prebiotic chemistry studies?. (2008). Orig Life Evol Biosph **38(6):**469-88. DOI: 10.1007/s11084-008-9150-5, PMID: 18925425

