## Supplementary material for "Coenzyme-Protein Interactions since Early Life"

**Supplementary Figures**

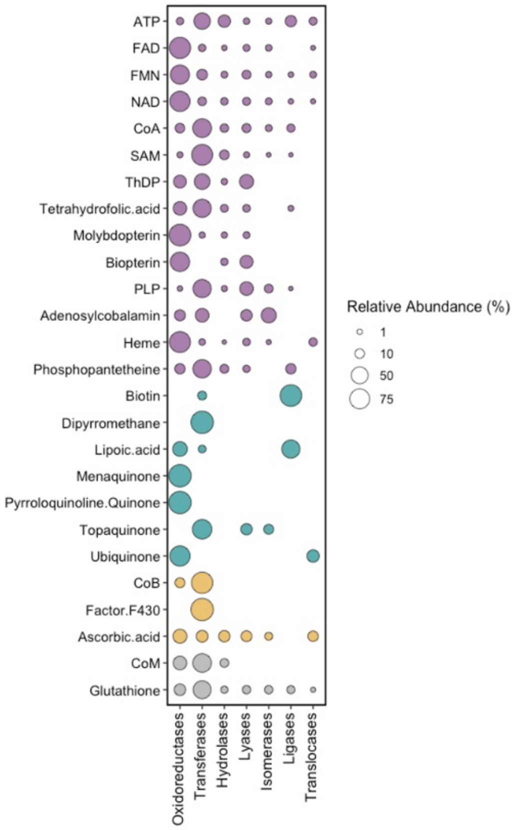


**Supplementary Figure 1:** Enzymatic class diversity per coenzyme class. All coenzymes are coloured by temporality: Ancient are shown in color purple, LUCA in Turquoise, Post-LUCA in yellow, and Unclassified cofactors in gray. The EC numbers were retrieved from SIFTS (Velankar et al. 2013; Dana et al. 2019).

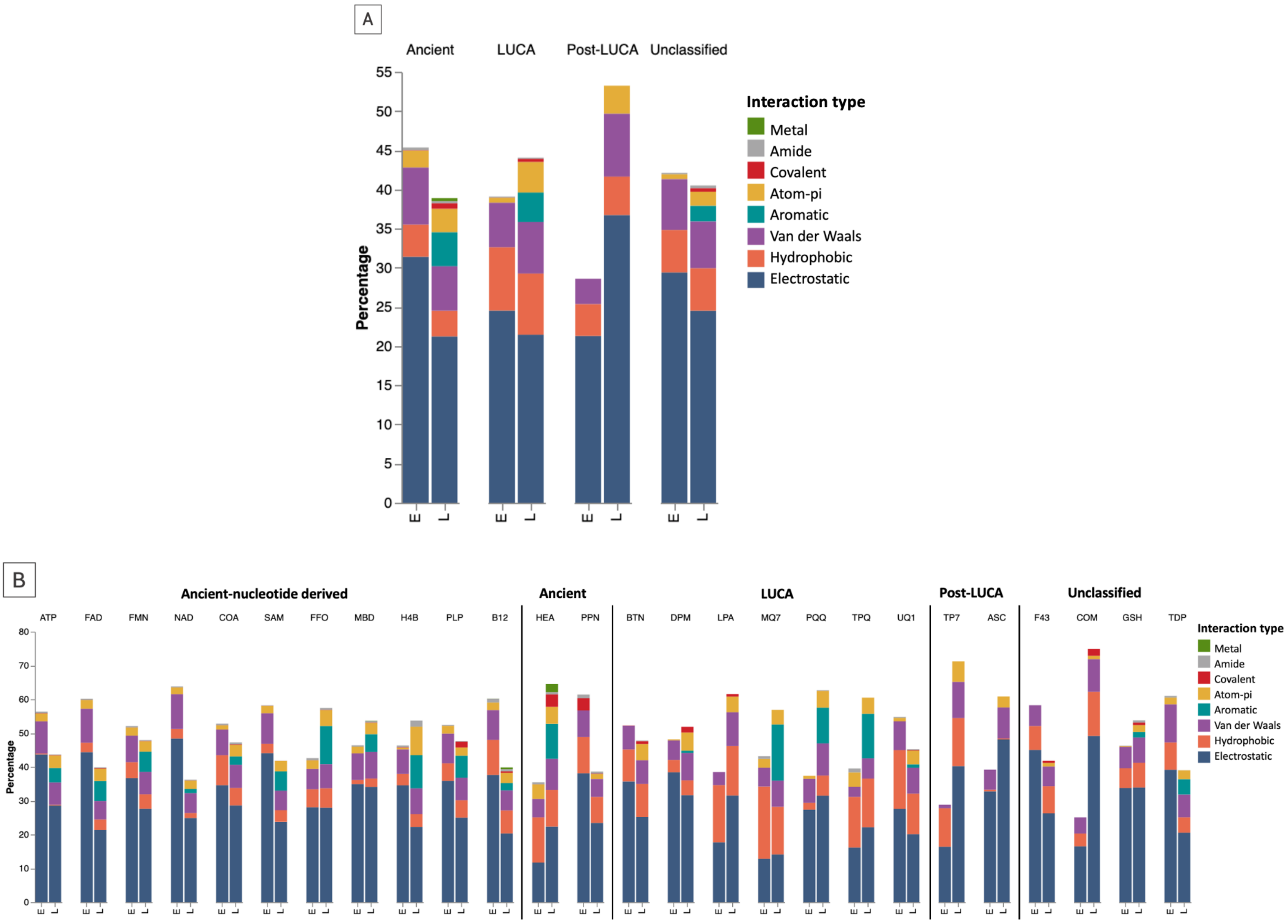


**Supplementary Figure 2:** Interaction types. (A) Interaction types for each amino acid-coenzyme binding event. (B) Interactions by coenzyme class. Early residues are shown as "E" and late as "L". The interactions were assigned by Arpeggio (Hubb et al. 2017).

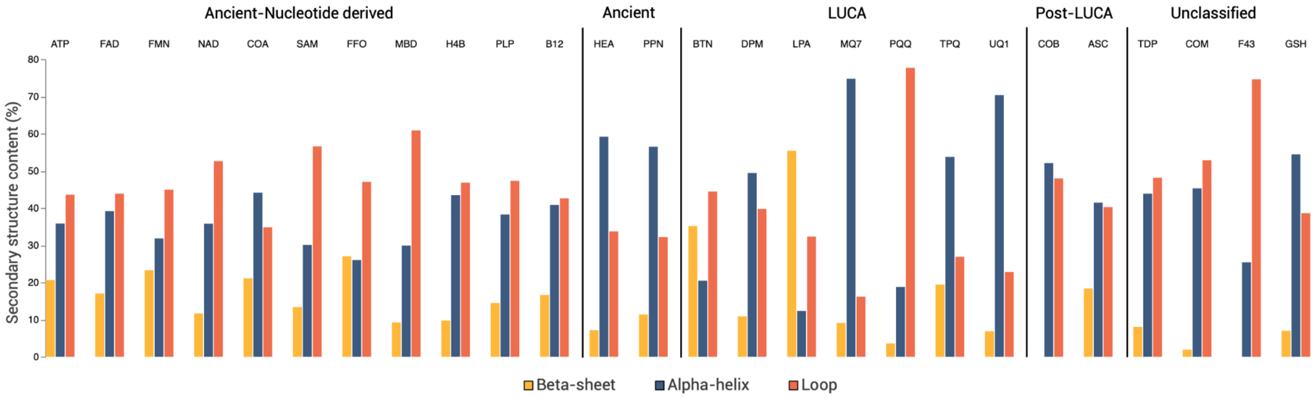


**Supplementary Figure 3:** Secondary structure content per cofactor class. All the structural assignments were obtained from SIFTS (Hubb et al. 2017).

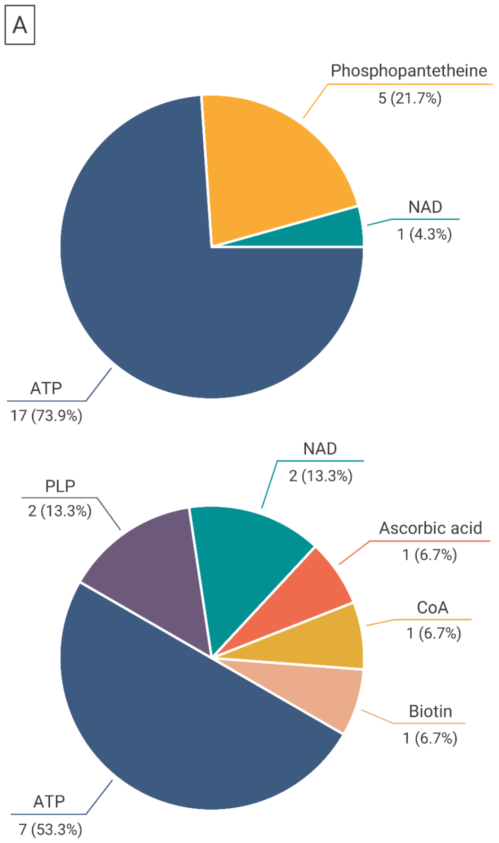


B

Coenzyme binding via only early amino acids

| Cofactor | Associated folds |
|---|---|
| ATP | Bacillus chorismate mutase-like<br>ATP_ligand,<br>Zincin-like |
| Phosphopantetheine | ACP-like |
| NAD | Not assigned |

Coenzyme binding via only late amino acids

| Cofactor | Associated folds |
|---|---|
| ATP | Retrovirus capsid protein<br>Ribonuclease H-like |
| PLP | Cradle loop barrel<br>Repetitive alpha hairpins |
| NAD | RNase A-like |
| Ascorbic acid | Repetitive alpha hairpins<br>Glycosyl hydrolase domain-like |
| CoA | Not assigned |
| Biotin | Not assigned |

**Supplementary Figure 4:** Cofactor binding with only early amino acids. ECOD X-group folds bound to those coenzymes are shown with the ConSurf (Ashkenazy et al., 2010; Ashkenazy et al., 2016) conservation scheme, and the enzyme cofactors are shown in spheres.

### Text files

**Supplementary file 1:** PDB codes assigned to each coenzyme class.

### Supplementary Tables

**Supplementary Table 1:** Identification of coenzymes in PDB.

**Supplementary Table 2:** Amino Acid Composition of the Coenzyme Binding Sites. Table 2A_90) Residue Composition at 90% Sequence Identity. Table 2B_30) Amino Acid Composition at 30% Sequence Identity.

**Supplementary Table 3:** Folds catalogue of ECOD X-groups in coenzyme binding sites.

**Supplementary Table 4:** Proteins and nucleic acids with coenzyme binding mediated by metallic ions and water molecules.

**Supplementary Table 5:** Amino acid fractional differences observed across all coenzyme binding sites.

**Supplementary Table 6:** Amino acid fractional differences observed across all non-phosphate containing coenzyme binding sites.

**Supplementary Table 7:** Coenzymes interacting with nucleic acids.

**Supplementary Table 8:** Chi-squared test of early versus late amino acid composition per coenzyme class.

**Supplementary Table 9:** Chi-squared test comparing early versus late residue composition across all coenzyme temporalities in different interaction types.

**Supplementary Table 10:** Average secondary structure content of the different coenzyme temporalities.
