## Supplementary Tables 5-10 for "Coenzyme-Protein Interactions since Early Life"

**Supplementary Table 5:** Amino acid fractional differences observed across all coenzyme binding sites. (A) Amino acid fractional difference of all coenzymes. (B) Amino acid fractional difference of all coenzymes at the residue level.

A

| Temporality | F (early) | F (late) | FDAA (F (early) - F (late)) |
|---|---|---|---|
| Ancient | 59.4 | 40.6 | 18.9 |
| LUCA | 52.3 | 47.7 | 4.7 |
| Post-LUCA | 37.1 | 62.9 | -25.9 |
| Unclassified | 53.2 | 46.8 | 6.4 |

B

| Amino acid | F (Ancient) | F (LUCA) | F (Post-LUCA) | F (Unclassified) | FDcofactor (F(Ancient) - F(LUCA)) | FDcofactor (F(Ancient) - F(Post-LUCA)) |
|---|---|---|---|---|---|---|
| Gly | 9.8 | 6.0 | 5.4 | 10.3 | 3.8 | 4.4 |
| Ala | 5.8 | 7.5 | 3.0 | 4.4 | -1.7 | 2.8 |
| Asp | 4.9 | 2.8 | 1.8 | 3.2 | 2.1 | 3.1 |
| Glu | 3.9 | 2.4 | 1.8 | 3.4 | 1.5 | 2.1 |
| Val | 5.7 | 5.4 | 9.2 | 6.4 | 0.3 | -3.5 |
| Ser | 7.6 | 5.8 | 3.3 | 6.7 | 1.8 | 4.3 |
| Ile | 5.1 | 6.2 | 4.4 | 4.6 | -1.2 | 0.6 |
| Leu | 7.5 | 9.1 | 3.4 | 6.5 | -1.6 | 4.1 |
| Pro | 3.1 | 1.5 | 2.4 | 3.7 | 1.5 | 0.7 |
| Thr | 6.3 | 5.6 | 2.5 | 4.1 | 0.6 | 3.8 |
| Lys | 4.0 | 3.2 | 5.9 | 2.0 | 0.8 | -1.9 |
| Phe | 5.6 | 6.7 | 16.6 | 9.4 | -1.1 | -11.0 |
| Arg | 6.0 | 8.7 | 10.2 | 5.8 | -2.6 | -4.2 |
| His | 4.6 | 4.6 | 7.8 | 5.1 | 0.0 | -3.2 |
| Asn | 4.3 | 3.9 | 5.7 | 2.5 | 0.4 | -1.5 |
| Gln | 2.9 | 3.0 | 3.6 | 5.3 | -0.1 | -0.7 |
| Cys | 2.0 | 3.7 | 0.9 | 3.2 | -1.7 | 1.1 |
| Tyr | 5.7 | 4.6 | 5.7 | 8.9 | 1.1 | 0.0 |
| Met | 3.1 | 3.0 | 5.1 | 3.2 | 0.0 | -2.1 |
| Trp | 2.5 | 6.4 | 1.5 | 1.3 | -3.9 | 1.0 |

**Supplementary Table 6:** Amino acid fractional differences observed across all non-phosphate containing coenzyme binding sites. (A) Amino acid fractional difference of non-phosphate containing coenzymes. (B) Amino acid fractional difference of non-phosphate containing coenzymes at residue level.

A

| Temporality | F (early) | F (late) | FDAA (F (early) - F (late)) |
|---|---|---|---|
| Ancient | 55.7 | 44.3 | 11.4 |
| LUCA | 52.3 | 47.7 | 4.7 |
| Post-LUCA | 40.2 | 59.8 | -19.5 |
| Unclassified | 48.7 | 51.3 | -2.6 |

B

| Amino acid | F (Ancient) | F (LUCA) | F (Post-LUCA) | F (Unclassified) | FDcofactor (F(Ancient) - F(LUCA)) | FDcofactor (F(Ancient) - F(Post-LUCA)) |
|---|---|---|---|---|---|---|
| Gly | 6.2 | 6.0 | 4.1 | 8.5 | 0.2 | 2.0 |
| Ala | 4.9 | 7.5 | 3.6 | 4.6 | -2.6 | 1.4 |
| Asp | 5.6 | 2.8 | 3.6 | 1.8 | 2.8 | 2.1 |
| Glu | 5.1 | 2.4 | 3.6 | 2.0 | 2.7 | 1.6 |
| Val | 5.6 | 5.4 | 3.6 | 7.1 | 0.3 | 2.1 |
| Ser | 5.4 | 5.8 | 6.5 | 7.0 | -0.4 | -1.1 |
| Ile | 6.0 | 6.2 | 4.7 | 4.0 | -0.3 | 1.2 |
| Leu | 9.5 | 9.1 | 1.8 | 6.3 | 0.4 | 7.7 |
| Pro | 2.8 | 1.5 | 4.7 | 3.8 | 1.3 | -1.9 |
| Thr | 4.5 | 5.6 | 4.1 | 3.7 | -1.1 | 0.4 |
| Lys | 2.5 | 3.2 | 5.9 | 2.3 | -0.7 | -3.4 |
| Phe | 7.9 | 6.7 | 10.1 | 11.2 | 1.2 | -2.1 |
| Arg | 5.4 | 8.7 | 8.9 | 6.9 | -3.2 | -3.4 |
| His | 6.2 | 4.6 | 10.7 | 4.3 | 1.7 | -4.4 |
| Asn | 3.0 | 3.9 | 6.5 | 1.4 | -0.9 | -3.5 |
| Gln | 2.3 | 3.0 | 7.1 | 6.3 | -0.7 | -4.8 |
| Cys | 2.6 | 3.7 | 1.8 | 4.1 | -1.1 | 0.8 |
| Tyr | 6.9 | 4.6 | 4.7 | 10.0 | 2.3 | 2.2 |
| Met | 3.4 | 3.0 | 1.2 | 3.4 | 0.4 | 2.2 |
| Trp | 4.0 | 6.4 | 3.0 | 1.5 | -2.4 | 1.0 |

**Supplementary Table 7:** Coenzymes interacting with nucleic acids.

| Coenzyme | Riboswitch | RNA aptamer | DNA aptamer |
|---|---|---|---|
| Adenosylcobalamin | 0 | 2 | 0 |
| ATP | 9 | 2 | 1 |
| Biopterin | 1 | 0 | 0 |
| Biotin | 0 | 2 | 0 |
| FMN | 12 | 1 | 0 |
| NAD | 8 | 0 | 0 |
| SAM | 41 | 0 | 0 |
| Tetrahydrofolic acid | 4 | 0 | 0 |
| ThDP | 6 | 0 | 0 |

**Supplementary Table 8:** *Chi-squared test* of early versus late amino acid composition per coenzyme class.

| Interaction type | Statistics |
|---|---|
| All | p-value = 0.0, chi-square = 2006.83, degrees of freedom = 26, critical value = 38.89. |

**Supplementary Table 9**: *Chi-squared test* comparing early versus late residue composition across all coenzyme temporalities in different interaction types.

| Interaction type | Statistics |
|---|---|
| Backbone | p-value = 0.01249, chi-square = 10.9, degrees of freedom = 3, critical value = 7.81 |
| Backbone & Side chain | p-value = 0.02875, chi-square = 9.04, degrees of freedom = 3, critical value = 7.81 |
| Side chain | p-value = 3.19E-26, chi-square =121.78, degrees of freedom = 3, critical value = 7.81 |

**Supplementary Table 10:** Average secondary structure content of the different coenzyme temporalities.

| Coenzyme/PDB | Secondary structure element | Mean | Standard deviation |
|---|---|---|---|
| PDB | beta | 15.024984 | 10.849963 |
| | loop | 33.100027 | 12.58042 |
| | helix | 51.874989 | 18.25092 |
| Ancient | beta | 15.024984 | 6.014811 |

| | | | |
|---|---|---|---|
| | loop | 33.100027 | 9.852286 |
| | helix | 51.874989 | 8.473325 |
| LUCA | beta | 20.037688 | 18.829335 |
| | helix | 42.819786 | 25.661259 |
| | loop | 37.142526 | 20.321961 |
| Post-LUCA | beta | 9.171598 | 12.970598 |
| | helix | 46.743117 | 7.527857 |
| | loop | 44.085285 | 5.442741 |
| Unclassified | beta | 4.218911 | 3.883016 |
| | helix | 42.233802 | 12.173157 |
| | loop | 53.547288 | 15.245032 |
